# The advantage of periodic over constant signalling in microRNA-mediated repression

**DOI:** 10.1101/2024.04.18.590057

**Authors:** Elsi Ferro, Candela L. Szischik, Alejandra C. Ventura, Carla Bosia

## Abstract

Cells have been found out to exploit oscillatory rather than constant gene expression to encode biological information. Temporal features of oscillations such as pulse frequency and amplitude have been shown determinant for the outcome of signaling pathways. However, little effort has been devoted to unveiling the role of pulsatility in the context of post-transcriptional gene regulation, where microRNAs (miRNAs) - repressors of gene expression - act by binding to RNAs. Here we study the effects of periodic against constant miRNA synthesis. We model periodic pulses of miRNA synthesis in a minimal miRNA-target RNA network by ODEs, and we compare the RNA repression to that resulting from constant synthesis of the repressor. We find that a pulsatile synthesis can induce more effective target RNA repression in the same timespan, despite an identical amount of repressor. In particular, a stronger fold repression is induced if the miRNA is synthesized at optimal frequencies, thereby showing a frequency preference behaviour - also known as “band-pass filtering”. Moreover, we show that the preference for specific input frequencies is determined by relative miRNA and target kinetic rates, thereby highlighting a potential mechanism of selective target regulation. Such ability to differentially regulate distinct targets might represent a functional advantage in post-transcriptional repression, where multiple competing targets are regulated by the same miRNA. Thereby analyzing a model with two RNA target species, we show how competition influences the frequency-dependent RNA repression. Eventually, we find that periodic miRNA expression can lead to exclusive frequency-dependent repression on distinct RNA species, and we show how this depends on their relative kinetics of interaction with the repressor. Our findings might have implications for experimental studies aimed at understanding how periodic patterns drive biological responses through miRNA-mediated signalling, and provide suggestions for validation in a synthetic miRNA-target network.

## 1. Introduction

A number of studies have highlighted that living cells are inherently dynamic: the concentrations and activities of many molecules previously assumed to stochastically fluctuate around fixed mean values have been shown as highly mutable over time. Live-cell time-lapse microscopy and fluorescent reporter genes have allowed to track the dynamic temporal behavior of proteins, thereby uncovering a picture where many regulators undergo pulses of activation and deactivation [1; 2]. Such pulses occur through temporal changes in their concentration or localization, on timescales that can span minutes or hours. How these oscillations are generated has been addressed by single-cell experiments, revealing that genetic circuits actively generate pulses of expression of key regulators and modulate pulse features such as frequencies and amplitudes [1]. Moreover, it has been shown that these features can determine the behavior of signaling pathways by driving crucial decision-making processes such as DNA repair, cell death and differentiation [1].

In parallel, theoretical studies have mathematically investigated the implications of pulsatility in signalling circuits and motivated its emergence in terms of biological functionality [3; 4]. Furthermore, they provided quantitative tools to predict biological outcomes based on temporal signal features. For instance, mathematical modelling predicted that simple circuits that involve binding and unbinding of two molecular species, such as ligand-receptor interaction, can behave as band-pass filters by selectively responding to certain frequencies of input signals [5; 6]. Nonetheless, periodic expression of regulator molecules has recently been observed also in the epigenetic context, as some post-transcriptional regulatory circuits were found to function in pulses: the interaction between miRNAs - short non-coding RNAs that repress gene expression - and their target genes was evidenced to shape gene expression oscillations [7; 8; 9; 10; 11].

MiRNAs are evolutionarily conserved RNAs with a length of 22-nt that act as posttranscriptional repressors of gene expression. Their repressive action is achieved by Watson-Crick base pairing with target RNAs, which can result in prevented translation for messenger RNAs or in an enhanced degradation [12]. Whether translation is affected or decay is increased depends on the strength of binding, that is the degree of miRNA-RNA complementarity [13; 14]. Since the condition for miRNA-RNA interaction is a binding of six consecutive nucleotides [15], a miRNA molecule can potentially interact with hundreds of different genes. Indeed, miRNA-mediated regulation is combinatorial, meaning that a single miRNA regulates multiple target genes, and a gene is typically targeted by several different miRNAs [13]. Thousands of mature miRNAs have been observed in humans, and miRNA-mediated circuits are increasingly being uncovered in key decision-making processes concerning development and differentiation [16; 17]. MiRNAs play fundamental roles in tumorigenesis, viral infection, and neurological diseases [18; 19; 20]. However, as the repressive action of miRNAs on target genes is weak compared to transcriptional regulation and rarely displays phenotypes [21], their pervasiveness is still partially unexplained. Theoretical and experimental studies suggested that their action might contribute to buffering gene expression noise, thereby working as fine tuners of genetic regulation [21; 22].

Periodic miRNA expression has been recently observed in diverse biological contexts ranging from development and differentiation to circadian rhythms [23; 24; 25; 26; 27]. Kim and co-workers showed that the let-7 miRNA can oscillate with a precise frequency to mask regulatory pulses of a transcription factor, thus maintaining a temporal gradient of its target functional for *C. Elegans* development [11]. Moreover, numerous miRNAs display circadian rhythmicity [28; 29; 30; 31].

However, the functional role of pulsatile post-transcriptional regulation is poorly understood - with theoretical predictions focusing mainly on small perturbations at the steady state [32] - and miRNAs have been largely overlooked by mathematical studies on periodic signalling. Therefore, we theoretically address pulsatile miRNA expression as opposed to constant expression, focusing on the functional role of oscillation frequency.

Using a minimal ODE model where a miRNA and an RNA species undergo simple binding and unbinding, we simulate pulsatile miRNA synthesis with variable frequency. We find that pulsatile miRNA synthesis can achieve a stronger average repression than a constant synthesis with identical miRNA-to-target dose in the same timespan. Furthermore, we show that the extent of repression depends on the frequency of pulses and thus uncover potential frequency preference behaviours. We then show that the advantage of pulsatile over constant synthesis and the system’s ability to be selective for specific frequency bands is determined by the relative miRNA and target kinetic features, suggesting that periodic miRNA signalling might provide a way to select targets with specific kinetics.

Since the ability to selectively regulate different targets can be particularly meaningful in the context of post-transcriptional repression, where many different RNA species are regulated by the same miRNA, we build a modified model where a miRNA targets two distinct RNA species. We thus show that the presence of a competitor can affect and shift the frequency-dependent repression of a target RNA. Eventually, we show that each competing target RNA can be selectively repressed in the same timespan using distinct miRNA synthesis frequencies. Our findings thus highlight periodic miRNA expression as a versatile mechanism of differential target regulation.

Overall, our findings suggest that periodicity might confer a functional advantage to post-transcriptional repression, and provide indications on experimental parameter tuning for validation in live cells.

## 2. Model M1

### 2.1. Modelling miRNA-target RNA interaction

The model was built using mass-action kinetics, as in [33]. Ordinary Differential Equations were non-dimensionalized by scaling variables and parameters with the RNA degradation rate constant and its synthesis rate constant (see Supplementary Material). As a results of this scaling, in the non-dimensional model one time unit corresponds to approximately 1/*ln*(2) ≈1, 44 RNA half-lives. This model considers a single target RNA species with one miRNA binding site. Both the RNA and the miRNA are synthesized through transcription. The two species can bind to form a complex, and each of them can be degraded both when unbound and when in complex. The model assumes that RNA and miRNA are degraded independently when in complex, as supported by previous data [34; 35; 36]: either the RNA is degraded and the miRNA is recycled into the system, or vice versa (Figure 1a,b).

**Figure 1:**
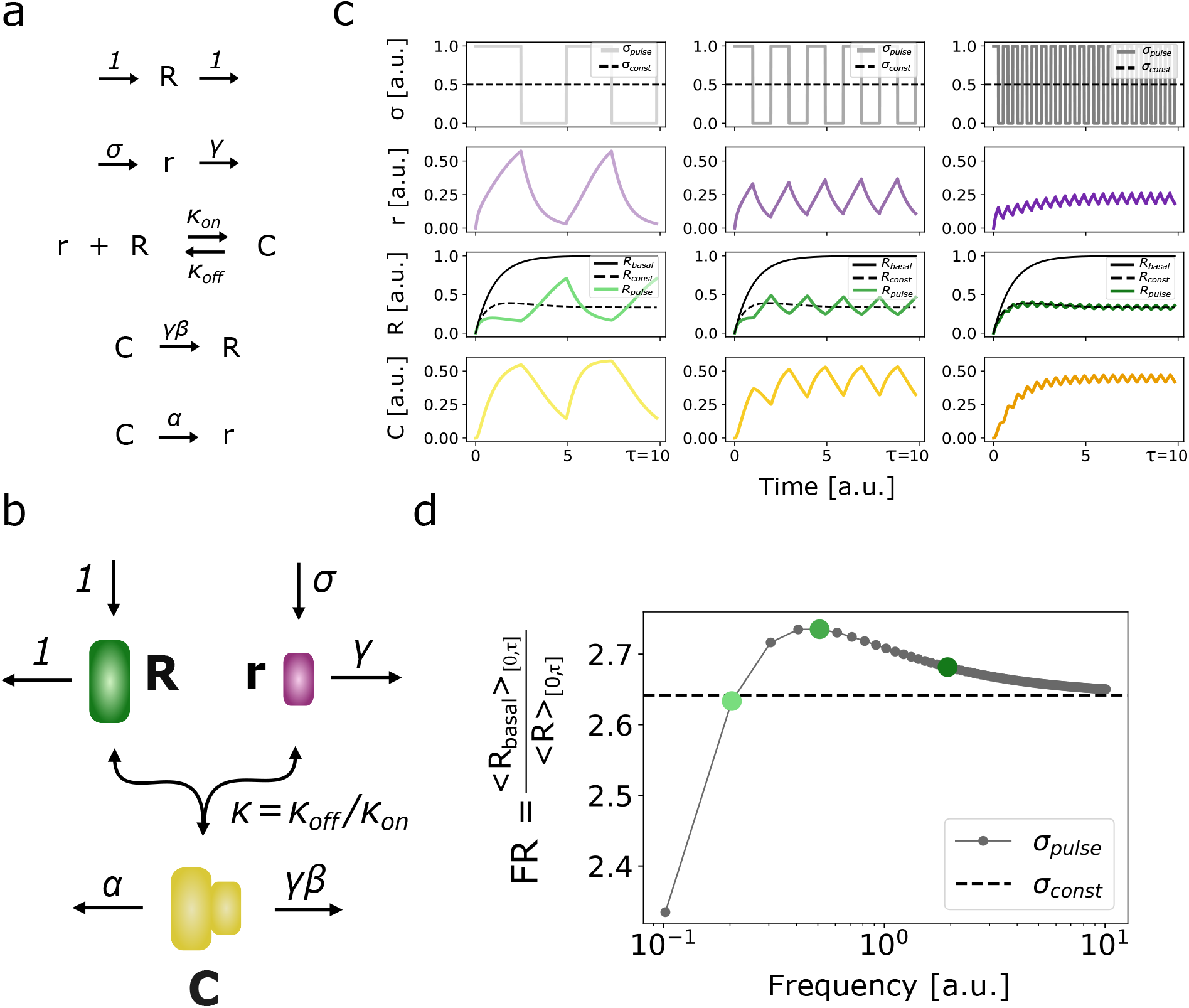
a) Schematic of biochemical reactions employed to model microRNA-RNA interaction. b) Schematic of microRNA-RNA interaction model. c) Example of molecular species temporal trajectories for three different input pulse frequencies. d) Example of fold repression computed in the [0, τ] time interval. Points represent FR values as a function of input miRNA synthesis pulse frequency (*FR*_*pulse*_(*f*)), with green dots marking the frequencies outlined in panel c. The black dashed line represents the FR value reached with a constant input of identical miRNA-to-RNA relative dose (*FR*_*const*_).

The dimensionless form of the mass-action model is described by the following ODE system:

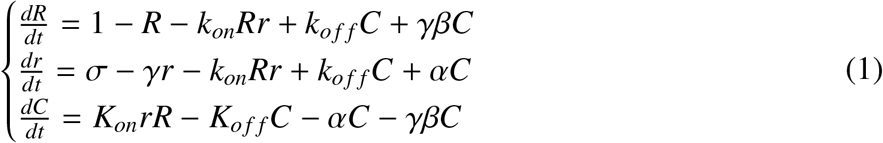

where *R, r* and *C* represent dimensionless concentrations of respectively unbound RNA, unbound miRNA and RNA-miRNA molecular complex. Due to the scaling choice (see Supplementary Material), the RNA production rate and the RNA degradation rate constant are scaled to 1. σ represents the scaled transcription rate constant of miRNA, and it is considered the signal input. *K*_*on*_ and *K*_*off*_ represent respectively the scaled association dissociation rates, thus yielding a scaled dissociation constant *K* = *K*_*off*_ /*K*_*on*_. γ describes the scaled degradation rate constant of miRNA relative to that of the RNA. α and β represent respectively the scaled degradation rates of RNA and miRNA in complex, that describe how fast each species is degraded in complex relative to its unbound form. These two parameters are particularly relevant, as miRNA-mediated regulation relies on RNA degradation upon binding [37], and similarly miRNA degradation can be affected by RNA binding [34].

### 2.2. Modelling periodic miRNA synthesis

Pulsatile miRNA expression is fed to the ODE systems by modelling the relative miRNA-RNA synthesis rate as a square pulse:

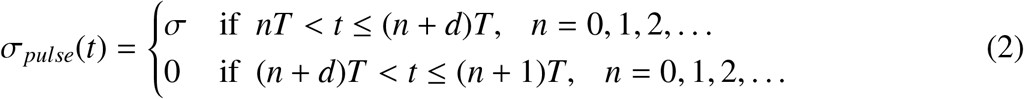

where *T* represents the pulse period and 0 < *d* < 1 represents its duty cycle.

Note that due to the scaling of time resulting from model nondimensionalization (see Supplementary Material), the pulse period *T* and thus its frequency *f* = 1/*T* are also scaled: a frequency *f* = 1 corresponds to pulses occurring every ≈ 1, 44 RNA half-lives - our nondimensional time unit.

### 2.3. Dose conservation in model M1

Considering the dimensional model M1, since the RNA is synthesized at a constant rate given by *S* _*R*_, its amount produced in a period *T* of the pulsatile input σ_*pulse*_ would be:

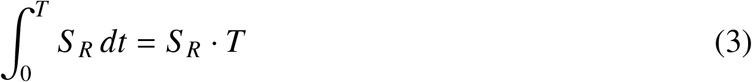

Conversely, the amount of miRNA produced in one period of σ_*pulse*_ would depend on the duty cycle *d* as:

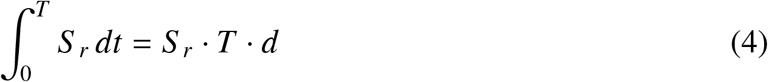

where *S* _*r*_ is the dimensional miRNA synthesis rate.

Therefore, considering that the scaled synthesis rate resulting from model nondimensionalization is 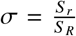 (see Supplementary Material), the dose of repressor synthesized relative to that of its target in a period will be:

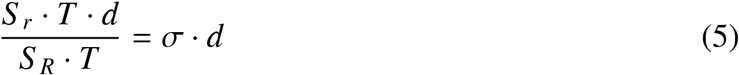

which results independent from the period size and thus from the input’s frequency. Thus, by keeping constant both the nondimensional synthesis rate σ and the duty cycle *d*, and feeding model M1 with an integer number of input pulses, we ensure that the amount of miRNA synthesized with respect to the RNA in the timespan of interest is maintained constant as we vary the input frequency. We call the quantity σ ·*d* “relative miRNA-RNA dose”. Despite alternative strategies of dose conservation rely on varying the duty cycle *d* and appropriately compensating the amplitude σ, we keep both values fixed as we vary the frequency. Indeed, we adopt a fixed duty cycle *d* = 0.5 relying on the observation that endogenous gene expression pulses commonly exhibit comparable ON and OFF pulse durations even beyond the context of circadian rhythms [38; 39].

Conversely, as the magnitude of gene expression pulses appears to be a feature characterizing the specific biochemical process, we still explore the role of pulse amplitude σ by parameter sensitivity analysis (see Supplementary Material).

### 2.4. M1 Model analysis

To analyze the model’s response, we use target gene fold repression as the primary output and we compute it as a function of input miRNA synthesis frequency. Fold repression is a commonly used measurement in the experimental quantification of miRNA-mediated repression, defined as the fold change between the average basal level of target gene expression and its average repressed level resulting from miRNA action:

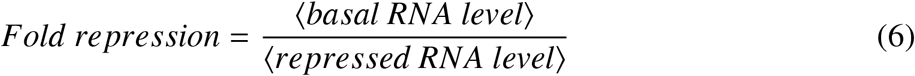

Since we aim at comparing the fold repression induced by periodic and constant miRNA syntheses, we compute the average fold repression achieved by both kinds of stimuli in an identical timespan. As such timespan we adopt the time needed by the constant input to induce steady state RNA level, which we call τ. We indeed consider τ - whose value is dependent on the system’s kinetic parameters - as a characteristic timescale required by the constantly stimulated system to reach its steady repression extent. In this way we address whether the average fold repression achieved by pulsatile stimuli of certain frequencies in the same timespan [0, τ] might overcome the one obtained by a constant stimulus with identical relative dose. We indicate as *FR*_*const*_ the fold repression value achieved by constant miRNA synthesis σ_*const*_:

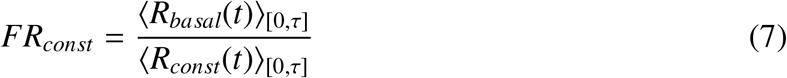

where *R*_*basal*_(*t*) represents the time-course of RNA achieved in the absence of miRNA and *R*_*const*_(*t*) represents the actual repressed RNA time-course induced by constant input synthesis σ_*const*_.

Similarly, we indicate as *FR*_*pulse*_ the fold repression value achieved by periodic miRNA synthesis σ_*pulse*_:

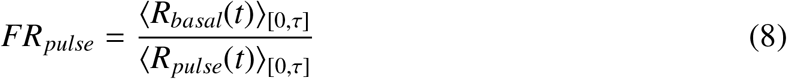

where *R*_*basal*_(*t*) represents again the RNA time-course achieved in the absence of miRNA and *R*_*pulse*_(*t*) represents the actual repressed RNA time-course induced by periodic input synthesis σ_*pulse*_. Note that both fold repression measurements - *FR*_*const*_ and *FR*_*pulse*_ - are ratios between variable values and are thus independent from the scaling of variables used for model nondimensionalization.

While the value of *FR*_*const*_ depends only on the system’s kinetic parameters, *FR*_*pulse*_ will depend also on the frequency of pulses and will be thus indicated as *FR*_*pulse*_(*f*).

To compute *FR*_*pulse*_(*f*) as a function of the frequency *f*, we numerically solve the ODE system for different frequencies of the input σ_*pulse*_. Since the relative miRNA-to-RNA dose depends only on the duty cycle *d* and the pulse amplitude σ as long as we consider only complete pulses, the considered time interval [0, τ] must contain an integer number of periods to ensure dose conservation, thus posing a constraint on the minimum frequency value that can be explored for a fixed value of τ: the lowest possible frequency will be *f*_*min*_ = 1/τ, whereas we fix the maximum frequency to *f*_*max*_ = 10^2^/τ.

Since we are interested in comparing outcomes of constant and pulsing inputs, we define a metric *A* that quantifies the repressive advantage of periodic over constant inputs as:

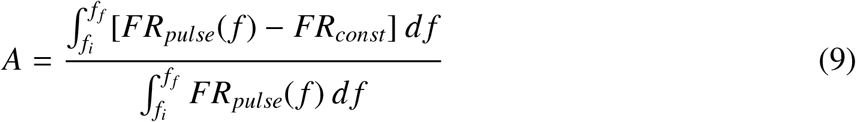

where *f*_*i*_ and *f* _*f*_ are respectively the lowest and the greatest frequency for which *FR*_*pulse*_(*f*) overcomes *FR*_*const*_. Note that *A* is bounded in the range [0, 1]. The broader the range of frequencies whose fold repression response *FR*_*pulse*_(*f*) overcomes that of the constant stimulus *FR*_*const*_ and the greater the difference between *FR*_*pulse*_(*f*) and *FR*_*const*_, the higher is the resulting value of *A*.

Since the frequency range where *FR*_*pulse*_(*f*) peaks may shift and/or narrow depending on the model’s kinetic parameters, we use a metric proposed by Romano and coworkers [40], that quantifies an observable’s selectivity *S* for a specific range of frequencies, given by:

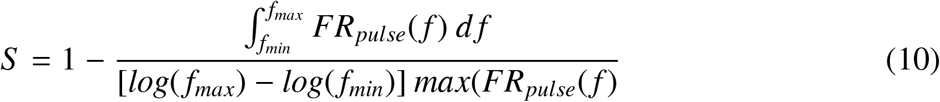

*S* is also bounded in the range [0, 1]. This metric considers the area and the height of the fold repression curve *FR*_*pulse*_ in relation to the frequency range covered, and thus measures the system’s frequency preference regardless of comparisons with the constant synthesis case.

Also the single frequency that allows to generate the highest value of *FR*_*pulse*_(*f*) - i.e. the system’s preferred frequency - may change depending on model parameters, and we thereby indicate it as:

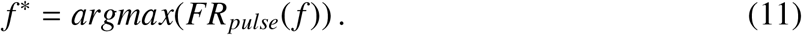

### 2.5. Periodic miRNA synthesis can achieve higher fold repression than constant synthesis in the characteristic timespan

To check whether any pulsing miRNA synthesis σ_*pulse*_ with a certain frequency could lead to stronger fold repression than a constant synthesis σ_*const*_ with identical relative dose σ/2, we chose a fixed parameter set with realistic kinetic rate values (see Materials and Methods), and we computed the average fold repression in the timespan [0, τ] both for the constant (*FR*_*const*_) and the periodic (*FR*_*pulse*_(*f*)) cases. In the periodic case, we numerically solved the system for all frequencies ranging from *f*_*min*_ = 1/τ to *f*_*max*_ = 10^2^/τ.

Interestingly, we found that some frequencies of periodic synthesis of the repressor lead to higher fold repression than constant synthesis (Figure 1c, d). As the frequency increases, the *FR*_*pulse*_(*f*) curve approaches *FR*_*const*_, showing that fold repression becomes progressively more similar to the one resulting from a constantly stimulated system.

### 2.6. Parameter sensitivity analysis of fold repression for constant and periodic miRNA synthesis in the characteristic timespan

To investigate how parameter variations impact the frequency preference of fold repression, 10^4^ parameter sets were randomly selected using Latin Hypercube sampling (LHS). Parameter distributions were estimated using values extracted from literature (see Supplementary Material), with the aim of covering a wide range of biologically plausible scenarios of RNA-miRNA interaction. Considering that due to non-dimensionalization the scaled parameters result as ratios of original dimensional parameters, a log-uniform sampling was used for each single scaled parameter. For each parameter set, we evaluated *FR*_*const*_ and *FR*_*pulse*_(*f*) in the range of frequencies that ensure relative dose conservation in the timespan [0, τ]. We subsequently computed the maximally preferred frequency *f* ^*^ and both the *FR*_*pulse*_(*f*) curve’s specificity *S* and the advantage *A*. Then, to determine which parameters most greatly impact each metric, we computed partial rank correlation coefficients (PRCC).

Which input frequency is the most advantageous for repression, i.e. *f* ^*^, resulted mostly determined by the relative miRNA-RNA degradation rate γ, showing that the reciprocal timescales of repressor and target decay are crucial in determining the preferred frequency band. Specifically, a fast repressor decay relative to the target seems associated to higher preferred frequencies, coherently with the fact that an unstable miRNA requires temporal accumulation to achieve its maximal repression.

The analysis also showed that the scaled RNA decay in complex α and the parameter representing relative miRNA/RNA degradation rate, γ, are the crucial parameters in determining the frequency selectivity of fold repression, together with the binding constant κ_*on*_ and the scaled miRNA decay in complex. Precisely, *S* resulted to be positively correlated with α, whereas negatively correlated with γ. We interpret the effect given by α as resulting from the fact that an enhanced target decay in complex amplifies the nonlinear effect given by miRNA-target interaction, thus resulting in an increased fold repression and a higher selectivity *S*. A similar and stronger correlation with α was found for the advantage *A*, as the amplification of repression given by the system’s nonlinearity becomes even more evident if one compares the outcomes of pulsatile and constant inputs. Oppositely, the negative coupling of both *A* and *S* with γ can be explained by the fact that an increased decay of the miRNA compared to its target leads to lower absolute fold repression values.

**Table 1:** Partial Rank Correlation Coefficients (PRCCs) between nondimensional model parameters and the advantage *A*, selectivity *S*, and the frequency of maximum fold repression *f* ^*^.

| | $\sigma$ | $k_{on}$ | $k_{off}$ | $\alpha$ | $\beta$ | $\gamma$ |
| --- | --- | --- | --- | --- | --- | --- |
| $A$ | 0, 10 | 0, 27 | -0, 14 | 0, 60 | -0, 24 | -0, 57 |
| $S$ | 0, 13 | 0, 25 | -0, 10 | 0, 38 | -0, 25 | -0, 27 |
| $f^*$ | 0, 24 | 0, 04 | 0, 04 | -0, 20 | 0, 09 | 0, 79 |

Note that the appearance of a preferred frequency in the *FR*_*pulse*_(*f*) curve is expected to be more frequent in the explored time interval [0, τ], corresponding to the transient state of the constantly stimulated system, than in a timespan corresponding to its steady state (e.g. [τ, 2τ]): the later the time window of observation, the more privileged the higher frequencies - which lead to a cumulative effect of miRNA action. Indeed, if we compute fold repression in the timespan [τ, 2τ] for random parameter sets identical to those used for computation in Figure 2a, we obtain a prevalence of high-pass filtering behaviors (see Supplementary Figure 12).

**Figure 2:**
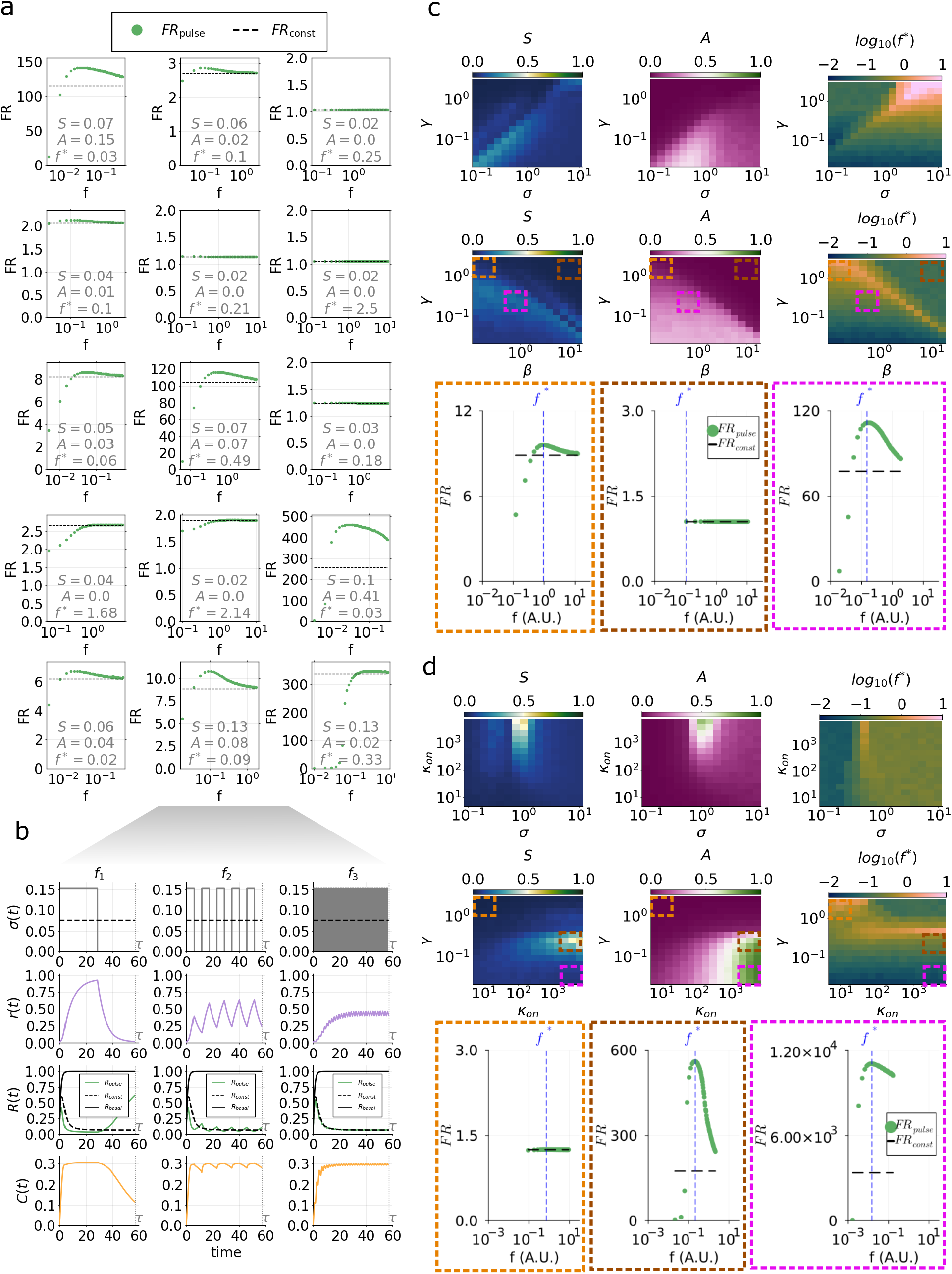
a) Random sampling of fold repression computed in the [0, τ] interval for either constant or periodic miRNA input synthesis. Green dots represent *FR*_*pulse*_(*f*) values, whereas black dashed lines represent *FR*_*const*_ values. Each subplot refers to a parameter set sampled randomly by Latin Hypercube sampling. For each case, advantage (*A*), selectivity (*S*) and preferred frequency (*f* ^*^) values are reported in the plot. b) Example of molecular species temporal trajectories for a case exhibiting fold repression advantage of periodic over constant input miRNA synthesis and frequency preference. Timecourses in the interval [0, τ] are reported for three different input miRNA synthesis frequencies. c) Fold repression advantage (left plot), selectivity (middle plot) and preferred frequency (right plot) as a function of synthesis and degradation parameter pairs. Top heatmaps represent the three metric quantities as functions of the input miRNA synthesis σ and the relative miRNA-RNA degradation rate γ; bottom heatmaps represent the three metric quantities as functions of the miRNA degradation rate in complex relative to its degradation alone, β, and the relative miRNA-RNA degradation rate γ. Underlying plots represent fold repression curves *FR*_*const*_ and *FR*_*pulse*_ corresponding to parameter values highlighted in heatmaps with dashed squares. d) Fold repression advantage (left plot), selectivity (middle plot) and preferred frequency (right plot) as a function of binding and synthesis/degradation parameter pairs. Top heatmaps represent the three metric quantities as functions of the input miRNA synthesis σ and the miRNA-RNA binding rate κ_*on*_; bottom heatmaps represent the three metric quantities as functions of the miRNA degradation rate in complex relative to its degradation alone, β, and the miRNA-RNA binding rate κ_*on*_. Underlying plots represent fold repression curves *FR*_*const*_ and *FR*_*pulse*_ corresponding to parameter values highlighted in heatmaps with dashed squares.

#### 2.6.1. Relative miRNA-RNA kinetics controls frequency preference of fold repression

To explore more in detail the role of influential parameters - identified by our sensitivity analysis - in determining the degree of preference for a specific range of frequencies, we computed the three metric quantities *A, S* and *f* ^*^ as functions of pairs of parameters, keeping the remaining parameters fixed to mean range values (Figure 2c, Supplementary Figures 6, 7, 8).

The variation of miRNA synthesis (i.e. σ) together with miRNA or RNA degradation timescales (i.e. α, γ, β) in pairs showed that a trade-off between these parameter values controls fold repression selectivity. For instance, maximal selectivity values are either found for concomitantly low or concomitantly high σ and γ values, whereas for simultaneously low (high) and high (low) values of γ and β. In parallel, *A* displays maximal values for concomitantly slow miRNA synthesis and degradation, i.e. low σ and γ, showing that a slowly but stably expressed miRNA leads to the highest advantage of periodic over constant repressor synthesis. Coherently, fast miRNA synthesis and degradation shift the preferred frequency *f* ^*^ to higher values, thus privileging scenarios where the miRNA accumulates in time rather than oscillating.

By simultaneously varying the miRNA-RNA binding rate κ_*on*_ and the miRNA synthesis rate σ we observed that both selectivity, advantage and preferred frequency *f* ^*^ display maximal values for a critical σ value, an effect that becomes more evident for high binding rate values (Figures 2c, Supplementary Figures 6, 7, 8). Instead, when varying the miRNA-RNA binding rate κ_*on*_ together with the relative miRNA-RNA decay rate γ, we found that only selectivity *S* displays a maximum at critical values of the degradation parameter, whereas the advantage *A* becomes maximal for concomitantly high binding κ_*on*_ and low miRNA decay γ rates. Note indeed that the fold repression given by periodic miRNA synthesis might present a high advantage over constant synthesis, but does not necessarily exhibit selectivity for a narrow range of frequencies. Interestingly, the maximal preferred frequency *f* ^*^ was found to switch at a critical value of κ_*on*_ from the highest to lower γ decay rate values, coherently with the fact that a higher miRNA-RNA binding affinity leads to accumulation of the repressor despite its reduced stability.

In general, we found that while the system’s synthesis and degradation parameters control frequency preference behaviours in a trade-off-like manner, their variation coupled with miRNA-RNA binding and unbinding parameters generates optimality in frequency preference.

### 2.7. Impact of initial miRNA and RNA conditions on advantage, selectivity and preferred frequency

Since the fold repression response might be affected also by initial species concentrations, we fixed model parameters to median values of log-uniform distributions in the estimated ranges and we evaluated both *FR*_*const*_ and *FR*_*pulse*_(*f*) starting from different miRNA and RNA concentrations. With this approach, we aim to consider different situations where either the repressor or the target or both start being expressed after a specific cellular signal and interacting with the cognate species. In particular, we focus on four different initial conditions where: (i) both the miRNA and the target RNA start being synthesized from zero: *R*_0_ = 0 and *r*_0_ = 0; (ii) miRNA synthesis starts from zero and acts on a previously unrepressed RNA target, i.e. the initial RNA concentration corresponds to the non-interacting RNA steady state which is always 1 due to nondimensionalization: *R*_0_ = 1 and *r*_0_ = 0; (iii) conversely, the initial miRNA concentration corresponds to its maximal steady state level in the absence of target interaction, i.e. σ/γ: *R*_0_ = 0 and *r*_0_ = σ/γ; iv) both the miRNA and the RNA are initially expressed to their non-interacting steady state levels, corresponding respectively to σ/γ and 1, and start interacting afterwards: *R*_0_ = 1 and *r*_0_ = σ/γ.

By varying initial miRNA and RNA conditions in the respective ranges *r*_0_ = [0, σ/γ] and *R*_0_ = [0, 1], we explore all the four situations.

In this way we found that both selectivity and advantage increase with the initial miRNA concentrations and decrease with that of the target RNA, consistent with a stronger miRNA-mediated repression (Figure 3). Similarly, the preferred frequency shifts to higher values for high initial miRNA and low initial RNA concentrations, thus privileging temporal miRNA accumulation rather than oscillation.

**Figure 3:**
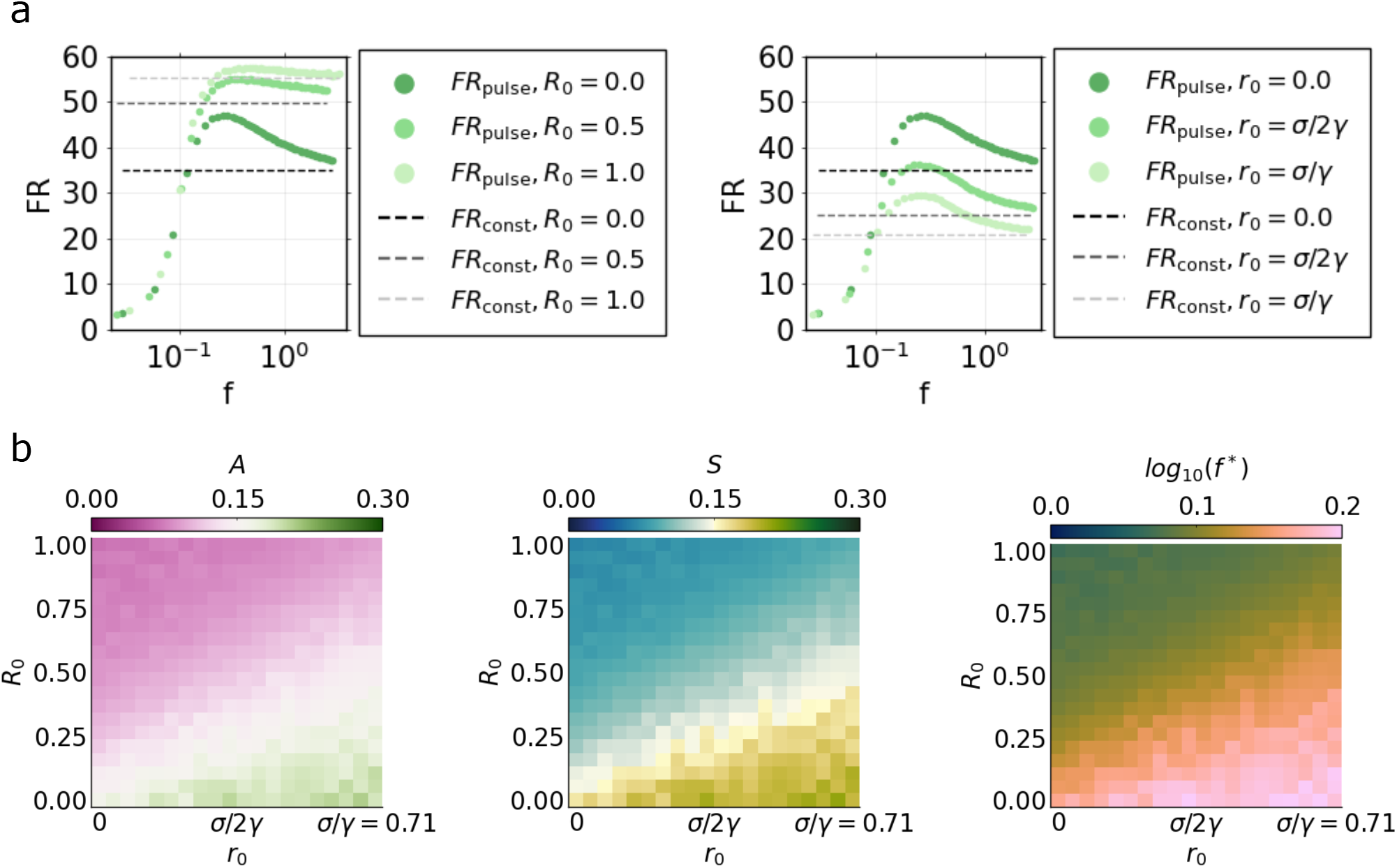
a) Fold repression computed in the [0, τ] interval by varying initial concentrations of miRNA or target RNA. Green dots represent *FR*_*pulse*_(*f*) values, whereas black dashed lines represent *FR*_*const*_ values. The left subplot represents *FR*_*pulse*_(*f*) and *FR*_*const*_ obtained for three different initial RNA concentrations (nondimensional values *R*_0_ = 0.0, 0.5, 1.0), maintaining null initial concentrations of miRNA and complex. The right subplot represents *FR*_*pulse*_(*f*) and *FR*_*const*_ obtained for three different initial miRNA concentrations (nondimensional values *R*_0_ = 0.0, σ/2γ = 0.36, σ/γ = 0.71), maintaining null initial concentrations of target RNA and complex. b) Advantage *A* (left panel), selectivity *S* (middle panel) and preferred frequency *f* ^*^ (right panel) as functions of initial miRNA and target RNA concentrations.

## 3. Model M2

M2 model is based on assumptions identical to model M1, but considers two target RNA species with one miRNA binding site each. Similarly to the first model, both the miRNA and the RNAs are synthesized through transcription and each RNA species can form a complex with the miRNA. All species can be degraded both when unbound and when in complex, with independent miRNA and RNA degradation similar to model M1 (Figure 4a, b).

**Figure 4:**
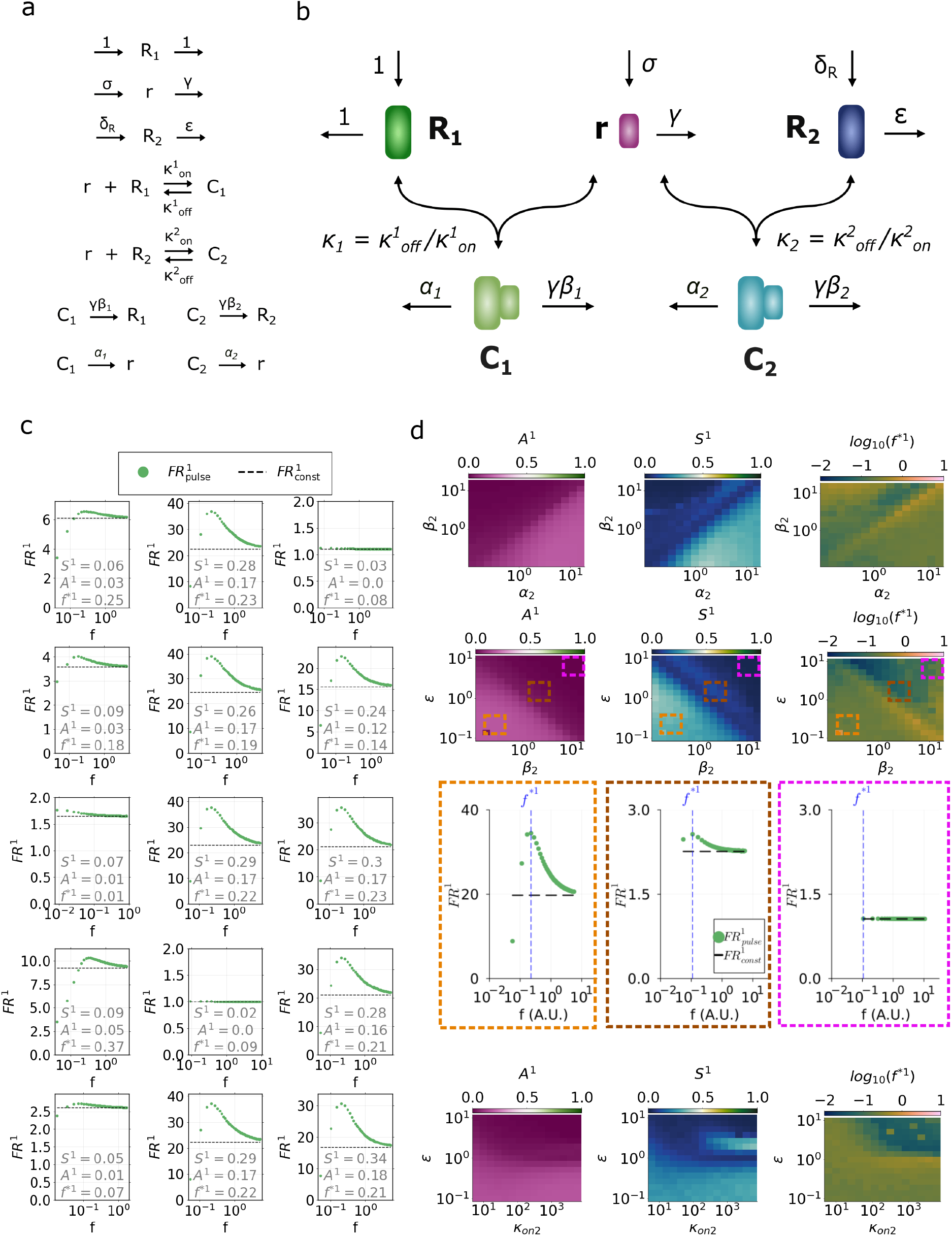
Analysis of fold repression given by periodic or constant miRNA synthesis in the presence of a competitor RNA target. a) Biochemical reactions considered in the nondimensional model M2. b) Schematic of nondimensional model M2. c) Random sampling of target *R*_1_ fold repression computed in the [0, τ_1_] interval for either constant or periodic miRNA input synthesis. Green dots represent 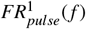. Black dashed lines represent 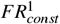. Each subplot refers to a set of parameters of the competitor target, i.e. *R*_2_, sampled randomly by Latin Hypercube sampling. For each case, values of target *R*_1_ advantage (*A*^1^) selectivity (*S* ^1^) and preferred frequency (*f* ^*1^) are reported in the plot. d) Fold repression advantage (left plot), selectivity (middle plot) and preferred frequency (right plot) as a function of synthesis and degradation parameter pairs. Top heatmaps represent the three metric quantities as functions of the miRNA and competitor RNA decay rates in complex, β_2_ and α_2_; the second row of heatmaps reports the three metric quantities as functions of the miRNA degradation rate in complex, β_2_, and the relative target degradation rate ϵ. Underlying plots represent fold repression curves 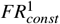 and 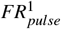 corresponding to parameter values highlighted in heatmaps with dashed squares. d) Bottom heatmaps represent the three metric quantities as functions of the miRNA-RNA binding rate κ_*on*_ and the relative target decay rate ϵ.

The dimensionless form of the mass-action model is described by the following ODE system:

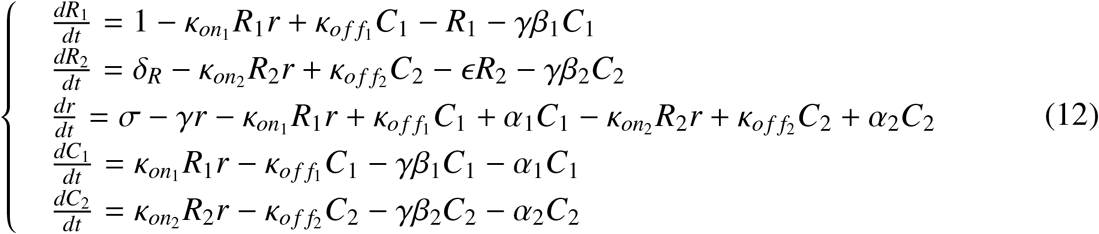

where *R*_1_ and *R*_2_ represent dimensionless concentrations of the two unbound RNA species, *r* is the dimensionless concentration of unbound miRNA, and *C*_1_ and *C*_2_ represent the two molecular complexes formed with the miRNA by the two distinct RNA species. Here, σ represents the synthesis rate constant of miRNA relative to that of the first RNA target. 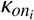 and 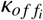 represent miRNA binding and unbinding rates of the *i*-th RNA species. α_*i*_ describes the degradation rate of the *i*-th RNA in complex relative to that of its unbound form. β_*i*_ is the degradation rate of miRNA in the *i*-th complex relative to its unbound form. γ is the degradation rate of unbound miRNA relative to that of the first RNA species. δ and ϵ describe respectively the synthesis and the degradation rate of the second RNA species with respect to the first.

### 3.1. Dose conservation in model M2

Considering the modified model M2 - where the miRNA targets two distinct RNA species - σ represents the synthesis rate of miRNA relative to the first RNA, and δ represents the synthesis rate of the second RNA species relative to the first. In this case, adopting a similar approach as for model M1, the dimensional amount of target 1 synthesized in one period of the periodic input would be:

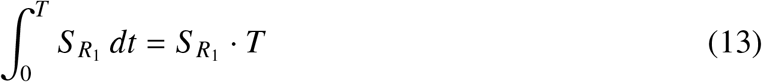

and similarly, the amount of target 2 synthesized in one period of the periodic input would be:

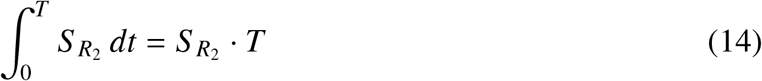

whereas the miRNA dose synthesized in one period would be again given by eq. (4). Therefore, the amount of repressor synthesized in one period relative to the first RNA species would result, similar to the M1 model case:

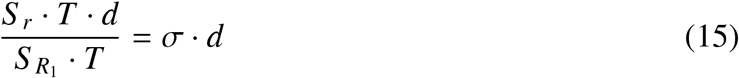

whereas its amount synthesized in one period relative to the second RNA species is:

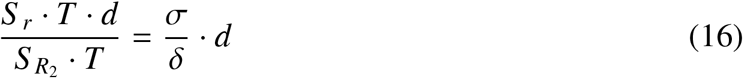

Thus, model M2 requires conserving both quantities σ· *d* and 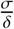 stant the amount of repressor synthesized relative to each target in a complete pulse. Therefore, since we assume a fixed duty cycle value of 0.5, we fix parameters σ and δ while varying the frequency of pulses. In the context of model M2 we will refer to the quantities σ · *d* and 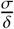 respectively as “relative miRNA-RNA1 dose” and “relative miRNA-RNA2 dose”.

### 3.2. M2 model analysis

To analyze the competitive model’s response, we use analogous fold repression measurements specific to each target, i.e. 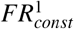 and 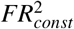 will refer to responses generated by constant miRNA synthesis while 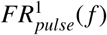 and 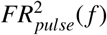 to responses generated by periodic miRNA synthesis relative to the first and second target RNA species.

We are interested in studying how the kinetics of a competitor RNA can affect the frequency-dependent behaviour of a target. Additionally, we aim to understand whether periodic miRNA expression may be able to tune the repression of different RNA targets to distinct pulse frequencies, and how the occurrence of this exclusive regulation scenario depends on the relative kinetics of competitors.

With the aim of comparing responses achieved by periodic and constant inputs in the characteristic timespan of one target, fold repression is always computed in the same timespan [0, τ_1_], which corresponds to the time needed by the constant stimulus to induce steady state of the first target.

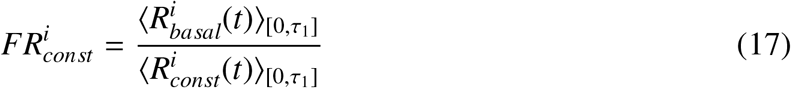

with the index *i* = 1, 2 referring either to the first or the second target.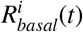 represents the RNA timecourses achieved in the absence of miRNA, 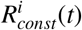 represents the actual repressed RNA timecourse induced by constant input synthesis σ_*const*_.

Analogously:

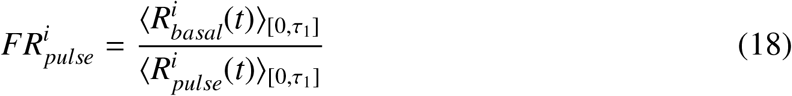

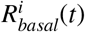 represents the RNA timecourses achieved in the absence of miRNA, and 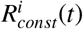 represents the actual repressed RNA timecourse induced by periodic input synthesis σ_*pulse*_.

Similarly to the M1 model, relative dose conservation is ensured regardless of frequency as long as the timespan [0, τ] contains an integer number of periods, and thus we use *f*_*min*_ = 1/τ_1_ and *f*_*max*_ = 10^2^/τ_1_ as lower and upper bounds for the explored frequency range.

We analogously compute the preferred frequencies for both RNA targets, *f* ^*1^ and *f* ^*2^, i.e. frequencies that generate the highest values of 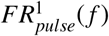 and 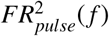 respectively, as:

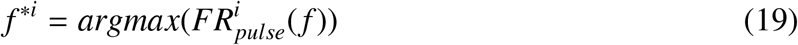

To evaluate the extent of frequency preference displayed by each target, we compute selectivities *S* _1_ and *S* _2_, which are always bounded in [0, 1] as:

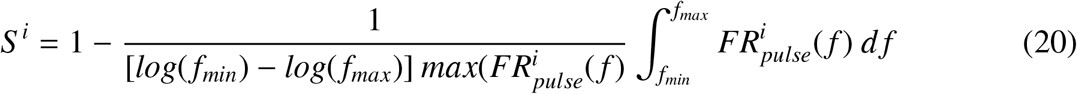

where *i* = 1, 2.

Analogously, we compute the fold repression advantage, bounded in [0, 1], displayed by each target upon pulsatile stimulation with respect to the constant input:

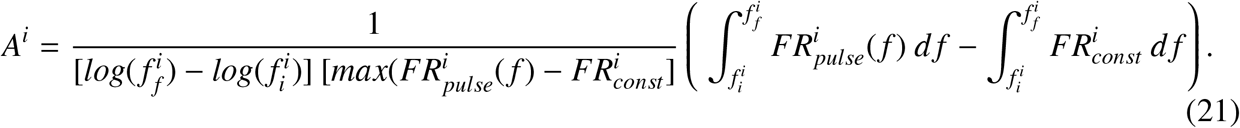

### 3.3. Sensitivity analysis of fold repression in the characteristic timespan to variation of competitor RNA kinetic parameters

To investigate how the addition of a competitor target affects the fold repression of the first target as a function of frequency, we performed a random parameter sampling using Latin Hypercube sampling. We fixed parameters relative to the first target, i.e. *R*_1_, and we randomly sampled parameters of the competitor, i.e.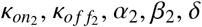, α_2_, β_2_, δ and ϵ. Fixed parameters correspond to median values of the log-uniformly distributed values in ranges estimated for model M2, whereas for the remaining parameters we drew 10^4^ sets from log-uniform distributions in the estimated ranges (see Supplementary Material).

First, we numerically solved the ODE system with constant miRNA synthesis rate σ_*const*_ to estimate the time τ_1_ needed by the first target RNA to reach steady state, using a relative tolerance threshold of 10^−6^. We thus defined the frequency values yielding an integer number of pulses in the range [0, τ_1_] to ensure relative dose conservation (see section 3.1). For each permitted frequency, we numerically solved the ODE system with periodic miRNA synthesis rate σ_*pulse*_, and we computed the average fold repression of target *R*_1_ as a function of frequency, i.e. 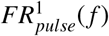,as indicated in equation 18.

Eventually, having the range of permitted frequencies [1/τ_1_, 10^2^/τ_1_], and the values of 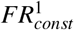 and 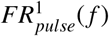 associated to each sampled parameter set of the competitor RNA, we calculated the three metric quantities for the *R*_1_ target, i.e. the maximally preferred frequency *f* ^*1^, the selectivity metric *S* ^1^ and the advantage metric *A*^1^, as described respectively in equations 19, 20, and 21.

The grid of plots in figure 4 shows examples of target *R*_1_ fold repression as a function of frequency for randomly selected parameter sets of the competitor target *R*_2_.

Then, to determine how the parameters of the competitor impact the frequency-dependent behavior of the first target, we calculated partial rank correlation coefficients (PRCC) of *f* ^*1^, *S* ^1^ and *A*^1^ with each sampled parameter. Correlation results are summarized in table 2.

**Table 2:**
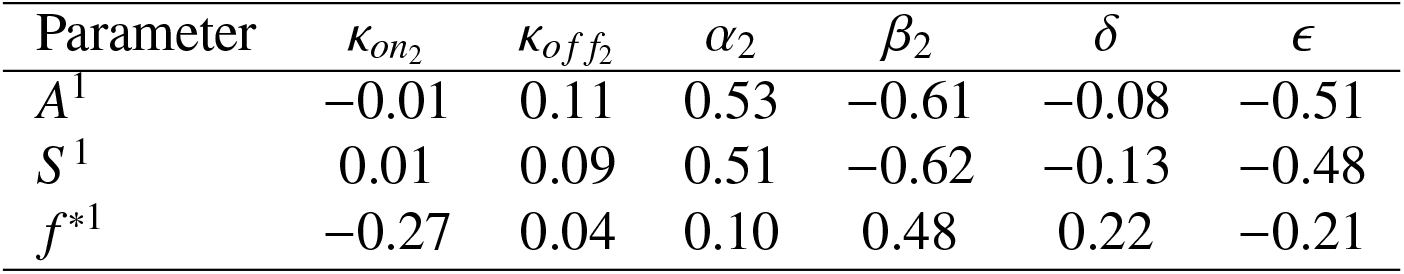
Partial rank correlation coefficients (PRCCs) between nondimensional model competitor’s parameters and the metric quantities relative to the first target *R*_1_: fold repression advantage (*A*^1^), fold repression selectivity (*S* ^1^), and frequency of maximum fold repression (*f* ^*1^).

In particular, we found that both *S* ^1^ and *A*^1^ are negatively correlated with β_2_ and positively with α_2_: a faster recycling of the competitor makes it dominant in the system, thus reducing the first target’s possibility to be repressed by the miRNA; whereas a faster recycling of the miRNA causes the opposite effect. Interestingly, both metrics are also anti-correlated with the parameter ϵ, showing that selectivity and advantage of *R*_1_ are enhanced by a longer half-life of the competitor. Note that since parameters of the first target are fixed to median range values, this effect might be relative only to this representative case of a typical target RNA, and might vary for a target with different kinetics. On the other hand, we found that the preferred frequency value of target *R*_1_ is strongly correlated with β_2_, δ and ϵ. Specifically, a faster degradation of the miRNA in complex with *R*_2_ (i.e. a higher value of β_2_) leads to a preference of the first target’s fold repression for higher frequencies. We can interpret this effect as due to the fact that a reduced availability of miRNA requires higher synthesis frequencies - and thus accumulation of the repressor - to achieve higher fold repression values. A similar result was obtained for δ, which controls the relative synthesis rate of the two competing targets: a faster synthesis of *R*_2_ causes a reduced availability of the miRNA to the first target, thus favoring again higher synthesis frequencies that lead to the repressor’s accumulation. Conversely, a faster decay of target *R*_2_ shifts the preference of *R*_1_ to lower frequencies of miRNA synthesis, as displayed by the negative correlation between ϵ and *f* ^*1^.

In general, these results suggest that whenever the presence of a competitor becomes dominant - leaving a restricted amount of repressor available - advantage and band-pass selectivity of the first target are shrinked and its preferred frequency is concomitantly shifted to higher values. The fact that the addition of a competitor with proper kinetics can modulate the preference of a target for a specific frequency might represent an interesting regulatory property of miRNA-mediated regulation.

### 3.4. Competitor RNA degradation and binding kinetics affects a target’s frequency preference of fold repression

To further explore the role of competitor RNA kinetic parameters in shaping a target’s degree of preference for a specific range of frequencies, we computed the selectivity *S* ^1^, the advantage *A*^1^ and the preferred frequency *f* ^*1^ as a function of pairs of parameters, keeping the remaining fixed to mean range values (Figures 4d and Supplementary Figures S4, S5, S6).

We found that a trade-off between distinct degradation timescales of the competitor RNA generates a switch from low to high advantage and selectivity of the first RNA target. For instance, a balance between a high miRNA recycling and a low enough competitor RNA recycling, α_2_ and β_2_, maintain *A*^1^ and *S* ^1^ in the high value region. Instead, a balance where both β_2_ and the competitor’s decay relative to the first target, ϵ, are low enough is necessary to maintain a high selectivity and advantage. Note that the preferred frequency of target 1, i.e. *f* ^*1^, presents a similar trade-off: the highest frequency values are found immediately outside of the high advantage and selectivity region, where the balance between competitor degradation parameters is lost.

If on the other hand we observe the role of binding/unbinding parameters of the competitor target, we find that if κ_*on*2_ is high enough, the fold repression of the first target becomes more selective - although not presenting the highest advantage - at a critical values of ϵ. This means that if the competitor RNA is slightly more unstable than the first, we achieve the highest *S* ^1^. Note that in this optimal selectivity region the system prefers lower frequency values. A similar optimality of *S* ^1^ is displayed at low κ_*o f f* 2_ values (Supplementary Figure S5).

Our results thus suggest that a competitor RNA’s half-life and its affinity to the repressor can be balanced to achieve an optimal frequency-driven repression on a first target RNA in its characteristic timespan.

### 3.5. Tuning miRNA-target RNA binding affinities and target RNA degradation can ensure differential regulation by desynchronization in frequency in the characteristic timespan

Eventually, to address the conditions where two competitor targets might show a preference for different input frequencies, we sampled the complete parameter space - including parameters of both targets - using Latin Hypercube sampling. Fixed parameters correspond to median values of the log-uniformly distributed values in ranges estimated for model M2 (sections 2 and 4), whereas for the remaining parameters we drew 10^4^ sets from log-uniform distributions in the estimated ranges (see Supplementary Material).

For each parameter set, we first numerically solved the ODE system with constant miRNA synthesis rate σ_*const*_ to estimate the time needed by the first RNA to reach the steady state, τ_1_, using a relative tolerance threshold of 10^−6^. We thus defined the frequency values yielding an integer number of pulses in the range [0, τ_1_] to ensure relative dose conservation (see section 3.1). For each frequency, we numerically solved the ODE system with periodic miRNA synthesis rate σ_*pulse*_, and we computed the average fold repression curves of both targets as a function of frequency, i.e. 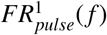 and 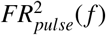, as indicated in equation 18, as well as the corresponding fold repression given by constant synthesis, 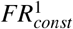 and 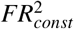 (see grid of plots in Figure 5a).

**Figure 5:**
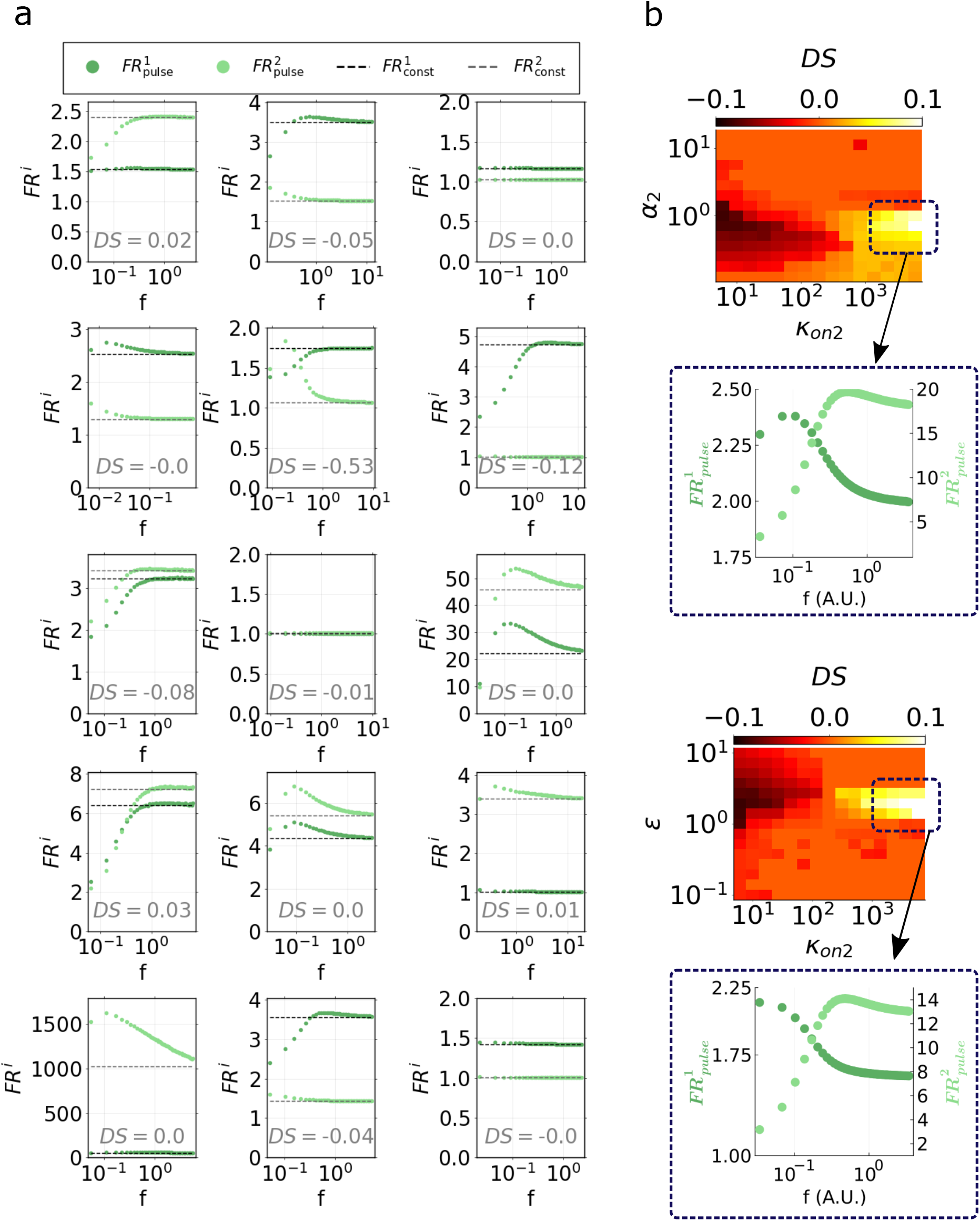
Analysis of differential target repression *DS*. a) Random sampling of fold repression of the two competing targets in the [0, τ_1_] interval for either constant or periodic miRNA input synthesis, computed sampling the whole parameter space of model M2. Dark and light green dots. represent respectively 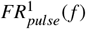 and 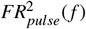. Black and grey dashed lines represent respectively 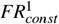 and 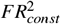 Each subplot refers to a different set of parameters. For each case, the value of differential selectivity *DS* is reported in the plot. b) Differential selectivity *DS* as a function of parameter pairs. The top heatmap represents *DS* as a function of the miRNA binding rate of the second target, κ_*on*2_, and the second target’s degradation rate in complex, α_2_; the underlying plot represents fold repression curves 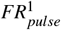 and 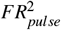 corresponding to parameter values highlighted in the heatmap with a dashed square. The bottom heatmap represents *DS* as a function of the miRNA-RNA binding rate κ_*on*2_ and the relative target decay rate ϵ. The underlying plot represents fold repression curves 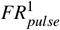 and 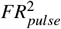 corresponding to parameter values highlighted in the heatmap with a dashed square.

Thus, to address situations where the fold repression of the two competing targets is maximal at different frequencies, we defined a metric to capture cases where the two targets’ selectivities *S* ^1^ and *S* ^2^ are concomitantly high and their preferred frequencies are distant, as:

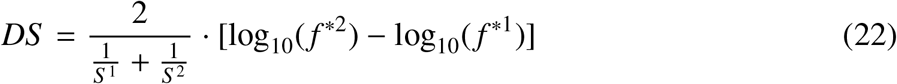

The harmonic mean of selectivities, given by 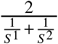, privileges cases where both *S* ^1^ and *S* ^2^ are similarly high, thus penalizing cases where only one of the two targets presents a high selectivity. Given the factor log_10_(*f* ^*2^) − log_10_(*f* ^*1^), we expect negative or positive *DS* values depending on the relative position in frequency of the maximal fold repression of the two target RNAs.

Thus, to investigate how the differential regulation of the two targets in frequency is affected by their relative kinetics, we calculated PRCCs (see results in Table 3) between our metric *DS* and the following parameter ratios: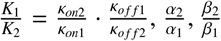, δ and ϵ.

**Table 3:** Partial rank correlation coefficients (PRCCs) between parameter ratios of the two target RNAs and the differential selectivity DS.

| Parameter | $K_2/K_1$ | $\alpha_2/\alpha_1$ | $\beta_2/\beta_1$ | $\delta$ | $\epsilon$ |
| --- | --- | --- | --- | --- | --- |
| $DS$ | 0.40 | -0.19 | -0.04 | 0.06 | -0.12 |

Interestingly, we found *DS* most strongly correlated with the relative dissociation constant of the two targets, *K*_2_/*K*_1_, showing that the separation of miRNA-target interaction timescales is important for the differential target repression controlled by frequency. Moreover, we found both the ratio between parameters α_2_ and α_1_ as well as the relative decay of the two targets, ϵ, slightly anticorrelated with *DS*.

This highlights the possibility of exclusive target regulation driven by periodic miRNA synthesis, and suggest that experimentally tunable kinetic rates - such as the relative miRNA-target affinity of the two competitor RNAs - can be adapted to achieve this exclusivity.

Given the role of relative target kinetics in determining the differential selectivity, we also explore the behaviour of such metric quantity, *DS*, varying the parameters that displayed the highest correlation. We thus compute *DS* as a function of κ_*on*2_ and α_2_, keeping the remaining parameters fixed, including the corresponding binding and unbinding rates of the first RNA target (Figure 5b). In this way we find that for either low or high affinity of the second target *R*_2_ with the repressor, κ_*on*2_, a narrow range of α_2_ values gives the highest occurrence of either negative or positive values of *DS* (Figure 5b, top heatmap). Similarly, by varying κ_*on*2_ and the relative decay rate of the two targets ϵ, we observe that in a narrow range of ϵ values the differential selectivity *DS* takes the highest absolute values, either negative or positive (Figure 5b, bottom heatmap). Note that this behaviour appears in a range of ϵ and α_2_ values where the competitor kinetics does not differ greatly from that of the first target. This suggests that small differences in the half-lives or in the decay induced by miRNA binding of the two targets can lead to their differential regulation in frequency, as long as their affinities to the repressor are dissimilar enough.

With these insights, we try to provide potential suggestions for an experimental tuning of target kinetics that might focus on the binding affinity between a miRNA and its targets and/or on the relative decay kinetics of the two competitors.

## 4. Discussion

The role of oscillatory gene expression has been extensively studied in the context of transcriptional regulation [1; 2], where periodic temporal concentrations of molecules have been shown to underlie the functionality of genetic pathways. Despite the tight connection between transcriptional and post-transcriptional regulatory interactions, the potential implications of periodic miRNA expression have been largely disregarded. Nevertheless, some experimental findings revealed the existence of periodically expressed miRNAs [23; 24; 25; 26; 27; 28; 29; 30; 31].

In this work we theoretically investigated the implications of periodic miRNA synthesis by focusing on repressive effects generated in the same timescale as constant synthesis. We consider a dynamical model wherein a microRNA binds a mRNA, resulting in a complex that can undergo unbinding or degradation through two pathways: either recycling the miRNA and degrading the mRNA or viceversa. We estimated the system’s timescale when fed with a constant synthesis rate as the time it takes mRNA to reach steady state. This enabled us to make a comparison between constant and periodic synthesis in inducing target RNA repression. With this approach, we could show that periodic miRNA synthesis can achieve a higher average fold repression than constant miRNA synthesis. According to our analysis, the maximal advantage of periodic over constant miRNA synthesis is found when the miRNA presence is overall stronger (i.e. longer half-life relative to the target, low degradation in complex, high initial concentration relative to the target). This is due to the inherent nonlinearity of the system, that amplifies the repression gap between periodic and constant miRNA synthesis. This suggests that revealing a repressive advantage of periodic over constant miRNA synthesis might be less challenging by combining stable miRNAs and unstable target RNAs, and thus privileging mRNAs over long-non-coding RNAs (lncRNAs) and circular RNAs (circRNAs) - which are known for their higher stability.

Interestingly, we showed that a high strength of binding between the miRNA and its target generates optimal advantage and frequency selectivity when the two are produced to equimolar concentrations (i.e. σ = 10^0^). This effect is due to the maximal nonlinearity of target response in this condition [41]. To experimental aims, this suggests that it might be possible to considerably strengthen the repression of a target RNA with high affinity to the miRNA by using periodic synthesis of the repressor - if their relative stoichiometries are properly tuned. We thus speculate that an artificial target with multiple miRNA-binding sites might enable detecting this effect.

We furthermore explored a few implications of competition between RNA targets for miRNA binding. By adding a competitor RNA with proper kinetics, we observed that a tuning of its kinetic rate constants can not only affect the repression of the first target driven by periodic miRNA synthesis, but most interestingly it can shift its preference for a specific range of frequencies. The idea that not only the single miRNA-target interaction but the entire competitive system including the multiple species at play might be kinetically tuned to achieve a precise target response is coherent with the well-known ceRNA hypothesis [42]. For instance, our predictions show that the frequency-dependent response of a target in the presence of a competitor mainly depends on their relative half-lives: this suggests that the observed concomitant targeting of RNA types that differ in their stability - such as circRNAs, lncRNAs and mRNAs - might represent a way to precisely tune repression driven by periodic miRNA expression. It is known for instance that both lncRNAs and circRNAs can serve as miRNA sponges that indirectly regulate miRNA target genes [43; 44]: according to our analysis, their action might be able to shift and/or sharpen the frequency-dependent response of a target in a way that depends on their degradation kinetics.

Eventually, our finding that a large gap between affinities of competing RNAs to the same miRNA can lead to their differential repression in the same timespan, highlights that proper input frequencies might enable the selective regulation of single targets despite the presence of multiple RNA species in the system. Again, miRNA-target binding kinetics emerges as a crucial element for the tuning of frequency-driven regulation. This scenario would be compatible with the fact that in principle a single miRNA can target RNA species with highly variable affinity - as the degree of complementarity is determined by the binding site sequence(s) located on the specific target transcript [12]. Moreover, the fact that moderate differences in target half-lives can bring out selective repression is compatible with differences in degradation kinetics detected among the main RNA types targeted by miRNAs (lncRNAs, circRNAs, mRNAs). For instance, the halflives of circRNAs were shown to be on average 2.5 times longer than those of mRNAs [45] - a difference compatible in our predictions both with the ability to shift the frequency preference of a competitor and with the possibility of differential regulation in frequency.

Our study is still limited by the use of a deterministic framework, which excludes the role of noise from our analyses - an element thought to be central in miRNA-mediated regulation. Given the nonlinearity of miRNA-target interaction, it was observed that extrinsic sources of noise can drive transitions between distinct states of target repression [46; 47]. The interplay between stochasticity and oscillation has attracted particular interest in the context of genetic regulation, where it has been predicted that the response of a nonlinear system to a weak periodic input signal can be amplified by the addition of optimal levels of noise. It is speculated that this phenomenon, also known as “stochastic resonance”, might be able to drive the switching between distinct states of gene expression [48; 49]. Interestingly, it was observed that the periodic binding and unbinding of a siRNA to mRNA in a live cell displays stochastic resonance [50]. Therefore, it would be of interest to investigate whether stochasticity might influence the repression of RNA targets driven by periodic miRNA synthesis in a biologically meaningful way.

## Author contributions

**CB** and **ACV** designed the project. **EF** and **CLS** performed all simulations, modeling analysis and writing. All authors have read and agreed on the final version of this article.

## Notes

### Competing Interest Statement

The authors have declared no competing interest.

## References

[1] Joe H. Levine, Yihan Lin, and Michael B. Elowitz. Functional Roles of Pulsing in Genetic Circuits. Science, 342(6163):1193–1200, 12 2013.

[2] Pawel Paszek, Dean A. Jackson, and Michael RH. White. Oscillatory control of signalling molecules. Current Opinion in Genetics & Development, 20:670–676, 2010.

[3] Jeremy E Purvis and Galit Lahav. Encoding and decoding cellular information through signaling dynamics. Cell, 152(5):945–56, 2 2013.

[4] Veena Venkatachalam, Ashwini Jambhekar, and Galit Lahav. Reading oscillatory instructions: How cells achieve time-dependent responses to oscillating transcription factors, 8 2022.

[5] Patrick A. Fletcher, Frédérique Clément, Alexandre Vidal, Joel Tabak, and Richard Bertram. Interpreting frequency responses to dose-conserved pulsatile input signals in simple cell signaling motifs. PLoS ONE, 9(4), 2014.

[6] P. Lánský, V. Křivan, and J. P. Rospars. Ligand-receptor interaction under periodic stimulation: A modeling study of concentration chemoreceptors. European Biophysics Journal, 30(2):110–120, 2001.

[7] Boyan Bonev, Peter Stanley, and Nancy Papalopulu. Microrna-9 modulates hes1 ultradian oscillations by forming a double-negative feedback loop. Cell reports, 2(1):10–18, 2012.

[8] Elizabeth S Fishman, Jisoo S Han, and Anna La Torre. Oscillatory behaviors of microrna networks: emerging roles in retinal development. Frontiers in Cell and Developmental Biology, 10:831750, 2022.

[9] Amitabha Nandi, Candida Vaz, Alok Bhattacharya, and Ramakrishna Ramaswamy. Mirna-regulated dynamics in circadian oscillator models. BMC systems biology, 3:1–16, 2009.

[10] Claude Gerard and Bela Novak. Microrna as a potential vector for the propagation of robustness in protein expression and oscillatory dynamics within a cerna network. PLoS One, 8(12):e83372, 2013.

[11] Dong Hyun Kim, Dominic Grün, and Alexander van Oudenaarden. Dampening of expression oscillations by synchronous regulation of a microrna and its target. Nature genetics, 45(11):1337–1344, 2013.

[12] Yimei Cai, Xiaomin Yu, Songnian Hu, and Jun Yu. A brief review on the mechanisms of mirna regulation. Genomics, Proteomics and Bioinformatics, 7(4):147–154, 2009.

[13] Zijun Luo, Xuping Xu, Peili Gu, David Lonard, Preethi H Gunaratne, Austin J Cooney, and Robert Azencott. Mirna regulatory circuits in es cells differentiation: A chemical kinetics modeling approach. PLoS One, 6(10):e23263, 2011.

[14] Xin Lai, Olaf Wolkenhauer, and Julio Vera. Understanding microrna-mediated gene regulatory networks through mathematical modelling. Nucleic acids research, 44(13):6019–6035, 2016.

[15] Daniel C Ellwanger, Florian A Büttner, Hans-Werner Mewes, and Volker Stümpflen. The sufficient minimal set of mirna seed types. Bioinformatics, 27(10):1346–1350, 2011.

[16] Himani Galagali and John K Kim. The multifaceted roles of micrornas in differentiation. Current opinion in cell biology, 67:118–140, 2020.

[17] Kathryn N Ivey and Deepak Srivastava. Micrornas as developmental regulators. Cold Spring Harbor perspectives in biology, 7(7):a008144, 2015.

[18] Beata Smolarz, Adam Durczyński, Hanna Romanowicz, Krzysztof Szyłło, and Piotr Hogendorf. Mirnas in cancer (review of literature). International journal of molecular sciences, 23(5):2805, 2022.

[19] Dan-Dan Cao, Lu Li, and Wai-Yee Chan. Micrornas: key regulators in the central nervous system and their implication in neurological diseases. International journal of molecular sciences, 17(6):842, 2016.

[20] Mona Fani, Milad Zandi, Majid Rezayi, Nastaran Khodadad, Hadis Langari, and Iraj Amiri. The role of micrornas in the viral infections. Current pharmaceutical design, 24(39):4659–4667, 2018.

[21] Jörn M Schmiedel, Sandy L Klemm, Yannan Zheng, Apratim Sahay, Nils Blüthgen, Debora S Marks, and Alexander van Oudenaarden. Microrna control of protein expression noise. Science, 348(6230):128–132, 2015.

[22] Héctor Herranz and Stephen M Cohen. Micrornas and gene regulatory networks: managing the impact of noise in biological systems. Genes & development, 24(13):1339–1344, 2010.

[23] Marc Goodfellow, Nicholas E Phillips, Cerys Manning, Tobias Galla, and Nancy Papalopulu. Microrna input into a neural ultradian oscillator controls emergence and timing of alternative cell states. Nature communications, 5(1):3399, 2014.

[24] Taeko Kobayashi and Ryoichiro Kageyama. Hes1 regulates embryonic stem cell differentiation by suppressing notch signaling. Genes to Cells, 15(7):689–698, 2010.

[25] Rongmin Chen, Matthew D’Alessandro, and Choogon Lee. Mirnas are required for generating a time delay critical for the circadian oscillator. Current Biology, 23(20):1959–1968, 2013.

[26] Xia Wang, Guihua Tian, Zhongfeng Li, and Lei Zheng. The crosstalk between mirna and mammalian circadian clock. Current Medicinal Chemistry, 22(13):1582–1588, 2015.

[27] Qian Gao, Lan Zhou, Su-Yu Yang, and Ji-Min Cao. A novel role of microrna 17-5p in the modulation of circadian rhythm. Scientific reports, 6(1):30070, 2016.

[28] Shunbin Xu, P Dane Witmer, Stephen Lumayag, Beatrix Kovacs, and David Valle. Microrna (mirna) transcriptome of mouse retina and identification of a sensory organ-specific mirna cluster. Journal of Biological Chemistry, 282(34):25053–25066, 2007.

[29] Hai Ying M. Cheng, Joseph W. Papp, Olga Varlamova, Heather Dziema, Brandon Russell, John P. Curfman, Takanobu Nakazawa, Kimiko Shimizu, Hitoshi Okamura, Soren Impey, and Karl Obrietan. MicroRNA Modulation of Circadian-Clock Period and Entrainment. Neuron, 54(5):813–829, 6 2007.

[30] Maocheng Yang, Jung-Eun Lee, Richard W Padgett, and Isaac Edery. Circadian regulation of a limited set of conserved micrornas in drosophila. BMC genomics, 9(1):1–11, 2008.

[31] Kyung-Ha Lee, Sung-Hoon Kim, Hwa-Rim Lee, Wanil Kim, Do-Yeon Kim, Jae-Cheon Shin, Seung-Hee Yoo, and Kyong-Tai Kim. Microrna-185 oscillation controls circadian amplitude of mouse cryptochrome 1 via translational regulation. Molecular biology of the cell, 24(14):2248–2255, 2013.

[32] Matteo Figliuzzi, Andrea De Martino, and Enzo Marinari. Rna-based regulation: dynamics and response to perturbations of competing rnas. Biophysical journal, 107(4):1011–1022, 2014.

[33] Benjamin Nordick, Polly Y. Yu, Guangyuan Liao, and Tian Hong. Nonmodular oscillator and switch based on RNA decay drive regeneration of multimodal gene expression. Nucleic Acids Research, 50(7):3693–3708, 2022.

[34] Manuel de la Mata, Dimos Gaidatzis, Mirela Vitanescu, Michael B Stadler, Corinna Wentzel, Peter Scheiffele, Witold Filipowicz, and Helge Großhans. Potent degradation of neuronal mi RNA s induced by highly complementary targets. EMBO reports, 16(4):500–511, 4 2015.

[35] Alessia Baccarini, Hemangini Chauhan, Thomas J Gardner, Anitha D Jayaprakash, Ravi Sachidanandam, and Brian D Brown. Kinetic analysis reveals the fate of a microrna following target regulation in mammalian cells. Current biology, 21(5):369–376, 2011.

[36] Francesco Ghini, Carmela Rubolino, Montserrat Climent, Ines Simeone, Matteo J Marzi, and Francesco Nicassio. Endogenous transcripts control mirna levels and activity in mammalian cells by target-directed mirna degradation. Nature communications, 9(1):3119, 2018.

[37] Stephen W. Eichhorn, Huili Guo, Sean E. McGeary, Ricard A. Rodriguez-Mias, Chanseok Shin, Daehyun Baek, Shu hao Hsu, Kalpana Ghoshal, Judit Villén, and David P. Bartel. MRNA Destabilization Is the dominant effect of mammalian microRNAs by the time substantial repression ensues. Molecular Cell, 56(1):104–115, 2014.

[38] Marie D Harton, Woo Seuk Koh, Amie D Bunker, Abhyudai Singh, and Eric Batchelor. p53 pulse modulation differentially regulates target gene promoters to regulate cell fate decisions. Molecular Systems Biology, 15(9):e8685, 2019.

[39] Hiromi Hirata, Shigeki Yoshiura, Toshiyuki Ohtsuka, Yasumasa Bessho, Takahiro Harada, Kenichi Yoshikawa, and Ryoichiro Kageyama. Oscillatory expression of the bhlh factor hes1 regulated by a negative feedback loop. Science, 298(5594):840–843, 2002.

[40] Daniel R. Romano, Matthew C. Pharris, Neal M. Patel, and Tamara L. Kinzer-Ursem. Competitive tuning: Competition’s role in setting the frequency-dependence of Ca2+-dependent proteins. PLoS Computational Biology, 13(11):1–26, 2017.

[41] Shankar Mukherji, Margaret S. Ebert, Grace X.Y. Zheng, John S. Tsang, Phillip A. Sharp, and Alexander Van Oudenaarden. MicroRNAs can generate thresholds in target gene expression. Nature Genetics, 43(9):854–859, 9 2011.

[42] Leonardo Salmena, Laura Poliseno, Yvonne Tay, Lev Kats, and Pier Paolo Pandolfi. A cerna hypothesis: the rosetta stone of a hidden rna language? cell, 146(3):353–358, 2011.

[43] Riccardo Taulli, Cristian Loretelli, and Pier Paolo Pandolfi. From pseudo-cernas to circ-cernas: a tale of cross-talk and competition. Nature structural & molecular biology, 20(5):541–543, 2013.

[44] Run-Wen Yao, Yang Wang, and Ling-Ling Chen. Cellular functions of long noncoding rnas. Nature cell biology, 21(5):542–551, 2019.

[45] Yehoshua Enuka, Mattia Lauriola, Morris E Feldman, Aldema Sas-Chen, Igor Ulitsky, and Yosef Yarden. Circular rnas are long-lived and display only minimal early alterations in response to a growth factor. Nucleic acids research, 44(3):1370–1383, 2016.

[46] Marco Del Giudice, Stefano Bo, Silvia Grigolon, and Carla Bosia. On the role of extrinsic noise in micrornamediated bimodal gene expression. PLOS Computational Biology, 14(4):1–26, 2018.

[47] Carla Bosia, Francesco Sgrò, Laura Conti, Carlo Baldassi, Davide Brusa, Federica Cavallo, Ferdinando Di Cunto, Emilia Turco, Andrea Pagnani, and Riccardo Zecchina. RNAs competing for microRNAs mutually influence their fluctuations in a highly non-linear microRNA-dependent manner in single cells, volume 18. 2017.

[48] Yong Xu, Juan Wu, Lin Du, and Hui Yang. Stochastic resonance in a genetic toggle model with harmonic excitation and lévy noise. Chaos, Solitons & Fractals, 92:91–100, 2016.

[49] Canjun Wang, Ming Yi, Keli Yang, and Lijian Yang. Time delay induced transition of gene switch and stochastic resonance in a genetic transcriptional regulatory model. BMC systems biology, 6:1–16, 2012.

[50] Shyamtanu Chattoraj, Shekhar Saha, Siddhartha Sankar Jana, and Kankan Bhattacharyya. Dynamics of gene silencing in a live cell: stochastic resonance. The Journal of Physical Chemistry Letters, 5(6):1012–1016, 2014.

[51] Lioudmila V. Sharova, Alexei A. Sharov, Timur Nedorezov, Yulan Piao, Nabeebi Shaik, and Minoru S.H. Ko. Database for mRNA half-life of 19 977 genes obtained by DNA microarray analysis of pluripotent and differentiating mouse embryonic stem cells. DNA Research, 16(1):45–58, 2 2009.

[52] Ron Milo, Paul Jorgensen, Uri Moran, Griffin Weber, and Michael Springer. BioNumbers The database of key numbers in molecular and cell biology. Nucleic Acids Research, 38(SUPPL.1), 10 2009.

[53] Cem Albayrak, Christian A. Jordi, Christoph Zechner, Jing Lin, Colette A. Bichsel, Mustafa Khammash, and Savaş Tay. Digital Quantification of Proteins and mRNA in Single Mammalian Cells. Molecular Cell, 61(6):914–924, 3 2016.

[54] Brian Reichholf, Veronika A. Herzog, Nina Fasching, Raphael A. Manzenreither, Ivica Sowemimo, and Stefan L. Ameres. Time-Resolved Small RNA Sequencing Unravels the Molecular Principles of MicroRNA Homeostasis. Molecular Cell, 75(4):756–768, 8 2019.

[55] Matteo J. Marzi, Francesco Ghini, Benedetta Cerruti, Stefano De Pretis, Paola Bonetti, Chiara Giacomelli, Marcin M. Gorski, Theresia Kress, Mattia Pelizzola, Heiko Muller, Bruno Amati, and Francesco Nicassio. Degradation dynamics of micrornas revealed by a novel pulse-chase approach. Genome Research, 26(4):554–565, 4 2016.

[56] William E. Salomon, Samson M. Jolly, Melissa J. Moore, Phillip D. Zamore, and Victor Serebrov. Single-Molecule Imaging Reveals that Argonaute Reshapes the Binding Properties of Its Nucleic Acid Guides. Cell, 162(1):84–95, 7 2015.

[57] Liang Meng Wee, C. Fabián Flores-Jasso, William E. Salomon, and Phillip D. Zamore. Argonaute divides Its RNA guide into domains with distinct functions and RNA-binding properties. Cell, 151(5):1055–1067, 11 2012.

