## supplemental text and figures for "The advantage of periodic over constant signalling in microRNA-mediated repression"

### Supplementary Material

#### 1. M1 model description and nondimensionalization

The eight chemical reactions included in our model are:

$$\varnothing \xrightarrow{s_R} \hat{R} \ , \ \hat{R} \xrightarrow{k_R^0} \varnothing \ , \ \varnothing \xrightarrow{s_r} \hat{r} \ , \ \hat{r} \xrightarrow{k_r^0} \varnothing \ , \ \hat{R} + \hat{r} \xrightarrow{k_{on}} \hat{C} \ , \ \hat{C} \xrightarrow{k_{off}} \hat{R} + \hat{r} \ , \ \hat{C} \xrightarrow{k_r^1} \hat{R} \ , \ \hat{C} \xrightarrow{k_R^1} \hat{r} \quad (1)$$

where $\hat{R}$, $\hat{r}$ and $\hat{C}$ represent respectively the RNA, the miRNA and the RNA-miRNA complex. The first two reactions represent respectively RNA synthesis and degradation, which occur with rate constants $S_R$ and $k_R^0$. Similarly, the third and fourth reactions describe miRNA synthesis and degradation along with their rates $S_r$ and $k_0^r$. The next two reactions represent miRNA-RNA binding and dissociation, whose constant rates are $k_{on}$ and $k_{off}$. The last two reactions describe respectively miRNA and RNA degradation when the two species are in complex, occurring at rates $k_r^1$ and $k_R^1$; these reactions imply that the species that does not undergo degradation - either the miRNA or the RNA - is recycled back into the system, and are thus often called *recycling reactions*.
Assuming the law of mass action, we describe the dynamics of the three molecule species by the following Ordinary Differential Equations (ODEs):

$$\begin{cases} \frac{d\hat{R}}{d\hat{t}} = S_R - k_{on}\hat{R}\hat{r} + k_{off}\hat{C} - k_R^0\hat{R} + k_r^1\hat{C} \\ \frac{d\hat{r}}{d\hat{t}} = s_r - k_{on}\hat{R}\hat{r} + k_{off}\hat{C} - k_r^0\hat{r} + k_R^1\hat{C} \\ \frac{d\hat{C}}{d\hat{t}} = k_{on}\hat{R}\hat{r} - k_{off}\hat{C} - k_R^1\hat{C} - k_r^1\hat{C} \end{cases} \quad (2)$$

To nondimensionalize the ODEs, we apply the following changes of variables:

$$\hat{t} = t/k_R^0 \ , \ \hat{R} = RS_R/k_R^0 \ , \ \hat{r} = rS_R/k_R^0 \ , \ \hat{C} = CS_R/k_R^0 \ . \quad (3)$$

Therefore, since the scaled time $t$ is proportional to the RNA half-life $t_{1/2} = \ln(2)/k_R^0$, one time unit in the nondimensional equations corresponds to $\hat{t} = \ln(2)t_{1/2} \approx 1.44\ t_{1/2}$. The nondimensionalized ODEs results as:

$$\begin{cases} \frac{dR}{dt} = 1 - \kappa_{on}Rr + \kappa_{off}C - R + \beta\gamma C \\ \frac{dr}{dt} = \sigma - \kappa_{on}Rr + \kappa_{off}C - \gamma r + \alpha C \\ \frac{dC}{dt} = \kappa_{on}Rr - \kappa_{off}C - \alpha C - \beta\gamma C \end{cases} \quad (4)$$

where:

$$\sigma = s_r/S_R \, , \, \gamma = k_r^0/k_R^0 \, , \, \kappa_{on} = \frac{k_{on}s_R}{{k_R^0}^2} \, , \, \kappa_{off} = \frac{k_{off}}{k_R^0} \, , \, \alpha = k_R^1/k_R^0 \, , \, \beta = k_r^1/k_r^0 \, .$$

Here, $\sigma$ represents the synthesis rate constant of miRNA relative to that of the RNA. $\gamma$ is the degradation rate constant of unbound miRNA relative to that of unbound RNA. $\alpha$ represents the degradation rate of RNA in complex relative to degradation in its unbound form. Analogously, $\beta$ represents the degradation rate of miRNA in complex relative to its unbound form.

#### 2. Estimation of biological rate constants

Prior to parameter sensitivity analysis of the fold repression response of the system, we estimated biologically meaningful ranges of parameters. Starting from the estimation of dimensional rate constants, we then derive ranges for nondimensional model parameters.
The median mammalian mRNA half-life in absence of post-transcriptional regulation, which corresponds to $k_R^0$ in our model, is estimated to be 4 hours [51]. The average diameter of a mammalian cell is $13\mu m$ [52], and thus the corresponding cell volume - considering the cell as a sphere - is $1.15 \times 10^{-10} L$. Since the number of mRNA molecules per gene in a single cell can range from a few copies to tens of thousands of copies, we adopt 200 molecules as mean value, in agreement with [47] and [53]. We can therefore calculate the mean mRNA molar concentration as $\bar{R} = n_R/N_A/V = 2.9 \times 10^{-10} M$, and estimate the RNA transcription rate as $s_R = k_R^0 \bar{R} = 1.4 \times 10^{-14} M s^{-1}$.
MiRNA half-lives are observed to be approximately four times the mRNA half-lives [54]. Therefore, we estimate the median of the scaled degradation rate $\gamma = \frac{K_r^0}{k_R^0}$ to be 1/4. However, as miRNA half lives can vary from about 4 hr up to 48 hr [55], it is possible that some miRNAs may have shorter half lives than their target RNAs, and we thus sample $\gamma$ log-uniformly in the range $[10^{-1}/4 - 10/4]$ as suggested by [33].
For the estimation of the miRNA-RNA association constant $k_{on}$ we adopt ranges spanning the orders of magnitude reported in [56]: $[10^6 - 10^9]$. Since the dissociation constant $K = k_{off}/k_{on}$ was estimated to be $3.7pM$ [57], we derive a suitable range for the miRNA-RNA dissociation rate constant as $k_{off}$ as $[3.7 \times 10^{-6}, 3.7 \times 10^{-3}] s^{-1}$. Then using the estimated values of $s_R$ and $k_R^0$ for scaling, we obtain ranges for nondimensional association and dissociation parameters: $\kappa_{on} = [6, 6 \times 10^3]$ and $\kappa_{off} = [7.7 \times 10^{-2}, 7.7 \times 1]$. In this way, using ranges for nondimensional parameters, we also explore scenarios where the number of RNA molecules per cell spans smaller and greater orders of magnitude with respect to the estimated mean value of 200 molecules.
Parameters $\alpha$ and $\beta$ were both sampled in the interval $[1/8, 16]$, estimated based on previous experimental data reported in [37; 34].
Eventually, to explore scenarios where miRNA synthesis is either faster, comparable or slower than synthesis of its target, we adopted $\sigma$ values ranging from $10^-1$ to $10^1$. Tables 4 and 5 report respectively estimated values of dimensional and nondimensional model parameters.

| Parameter | Biological meaning | Estimated median values / ranges |
|---|---|---|
| $k_R^0$ | RNA degradation rate | $4.8 \times 10^{-5} s^{-1}$ |
| $S_R$ | RNA transcription rate | $1.4 \times 10^{-14} M s^{-1}$ |
| $K$ | Dissociation constant | 3.7 pM |
| $k_{on}$ | Binding rate | $[10^6 - 10^9]\ M^{-1} s^{-1}$ |
| $k_{off}$ | Unbinding rate | $[3.7 \times 10^{-6} - 3.7 \times 10^{-3}]\ s^{-1}$ |

Table 4: Dimensional M1 model's parameter ranges.

#### 3. M2 model description and nondimensionalization

In this modified model we add a second RNA species able to bind the same miRNA. We thus consider one miRNA species and two RNA species that form distinct molecular complexes with the miRNA. The chemical reactions involved in this model are the following:

| Nondimensional parameter | Biological meaning | Range |
|---|---|---|
| $\sigma$ | Scaled miRNA transcription rate | $[10^{-1}, 10^{1}]$ |
| $\kappa_{on}$ | Scaled binding rate | $[6, 6 \times 10^{3}]$ |
| $\kappa_{off}$ | Scaled unbinding rate | $[7.7 \times 10^{-2}, 7.7 \times 10^{1}]$ |
| $\alpha$ | Bound relative to unbound RNA degradation rate | [1/8, 16] |
| $\beta$ | Bound relative to unbound miRNA degradation rate | [1/8, 16] |
| $\gamma$ | Scaled miRNA degradation rate | $[10^{-1}/4, 10/4]$ |

Table 5: Nondimensional M1 model's parameter ranges.

$$\varnothing \overset{s_{R_1}}{\rightarrow} \hat{R}_1 \quad \hat{R}_1 \overset{k^0_{R_1}}{\rightarrow} \varnothing \quad \varnothing \overset{s_{R_2}}{\rightarrow} \hat{R}_2 \quad \hat{R}_2 \overset{k^0_{R_2}}{\rightarrow} \varnothing \quad \varnothing \overset{s_r}{\rightarrow} \hat{r} \quad \hat{r} \overset{k^0_r}{\rightarrow} \varnothing \quad \hat{R}_1 + \hat{r} \overset{k_{on_1}}{\rightarrow} \hat{C} \quad \hat{R}_2 + \hat{r} \overset{k_{on_2}}{\rightarrow} \hat{C} \tag{5}$$

$$\hat{C} \overset{k_{off_1}}{\rightarrow} \hat{R}_1 + \hat{r} \quad \hat{C} \overset{k_{off_2}}{\rightarrow} \hat{R}_2 + \hat{r} \quad \hat{C} \overset{k^1_{r_1}}{\rightarrow} \hat{R}_1 \quad \hat{C} \overset{k^1_{r_2}}{\rightarrow} \hat{R}_2 \quad \hat{C} \overset{k^1_{R_1}}{\rightarrow} \hat{r} \quad \hat{C} \overset{k^1_{R_2}}{\rightarrow} \hat{r}$$

where $\hat{R}_1$, $\hat{R}_2$, $\hat{r}$, $\hat{C}_1$ and $\hat{C}_2$ represent respectively the two RNA species, the miRNA and the two RNA-miRNA complex species.
The first two reactions represent the synthesis and degradation of the first RNA species, which occur respectively at rates $s_{R_1}$ and $k^0_{R_1}$. Similarly, the next two reactions describe the synthesis and degradation of the second target, associated with rates $s_{R_2}$ and $k^0_{R_2}$. The fifth and sixth reactions represent miRNA synthesis and degradation along with rates $s_r$ and $k^0_r$. Next, we report reactions of binding and unbinding for both RNA targets, occurring respectively with rates $k_{on_1}$ and $k_{off_1}$, and $k_{on_2}$ and $k_{off_2}$. Eventually, reactions of miRNA degradation in complex (i.e. recycling of one of the two targets) occur with rates $k^1_{r_1}$ and $k^1_{r_2}$ for the two complex species, whereas those of target degradation in complex (i.e. miRNA recycling) follow respective rates $k^1_{R_1}$ and $k^1_{R_2}$.
Considering the law of mass action, we describe the dynamics of the five molecule species using the following Ordinary Differential Equations (ODEs):

$$\begin{cases} \frac{d\hat{R}_1}{d\hat{t}} = s_{R_1} - k_{on_1}\hat{R}\hat{r} + k_{off_1}\hat{C}_1 - k^0_{R_1}\hat{R}_1 + k^1_{r_1}\hat{C}_1 \\ \frac{d\hat{R}_2}{d\hat{t}} = s_{R_2} - k_{on_2}\hat{R}\hat{r} + k_{off_2}\hat{C}_2 - k^0_{R_2}\hat{R}_2 + k^1_{r_2}\hat{C}_2 \\ \frac{d\hat{r}}{d\hat{t}} = s_r - k_{on_1}\hat{R}\hat{r} - k_{on_2}\hat{R}\hat{r} + k_{off_1}\hat{C}_1 + k_{off_2}\hat{C}_2 - k^0_r\hat{r} + k^1_{R_1}\hat{C}_1 + k^1_{R_2}\hat{C}_2 \\ \frac{d\hat{C}_1}{d\hat{t}} = k_{on_1}\hat{R}_1\hat{r} - k_{off_1}\hat{C}_1 - k^1_{R_1}\hat{C} - k^1_{r_1}\hat{C}_1 \\ \frac{d\hat{C}_2}{d\hat{t}} = k_{on_2}\hat{R}_2\hat{r} - k_{off_2}\hat{C}_2 - k^1_{R_2}\hat{C} - k^1_{r_2}\hat{C}_2 \end{cases} \tag{6}$$

where $\hat{R}_1$ and $\hat{R}_2$ represent concentrations of the two unbound RNA species, $\hat{r}$ describes miRNA concentration, whereas $\hat{C}_1$ and $\hat{C}_2$ describe concentrations of the two RNA-miRNA complex species.
Adopting the same approach as for the M1 model, we make the following changes of variables using the synthesis and degradation rates of the first RNA, $s_{R_1}$ and $k^0_{R_1}$:

$$\hat{t} = t/k^0_R, \quad \hat{R}_1 = R_1 s_{R_1}/k^0_{R_1}, \quad \hat{R}_2 = R_2 s_{R_1}/k^0_{R_1}, \quad \hat{r} = r s_{R_1}/k^0_{R_1}, \quad \hat{C}_1 = C_1 s_{R_1}/k^0_{R_1}, \quad \hat{C}_2 = C_2 s_{R_1}/k^0_{R_1} \tag{7}$$

which yield the non-dimensional ODE system:

$$\begin{cases} \frac{dR_1}{dt} = 1 - \kappa_{on_1} R_1 r + \kappa_{off_1} C_1 - R_1 - \gamma\beta_1 C_1 \\ \frac{dR_2}{dt} = \delta_R - \kappa_{on_2} R_2 r + \kappa_{off_2} C_2 - \epsilon R_2 - \gamma\beta_2 C_2 \\ \frac{dr}{dt} = \sigma - \kappa_{on_1} R_1 r + \kappa_{off_1} C_1 - \gamma r + \alpha_1 C_1 - \kappa_{on_2} R_2 r + \kappa_{off_2} C_2 + \alpha_2 C_2 \\ \frac{dC_1}{dt} = \kappa_{on_1} R_1 r - \kappa_{off_1} C_1 - \gamma\beta_1 C_1 - \alpha_1 C_1 \\ \frac{dC_2}{dt} = \kappa_{on_2} R_2 r - \kappa_{off_2} C_2 - \gamma\beta_2 C_2 - \alpha_2 C_2 \end{cases} \tag{8}$$

where:

$$\sigma = s_r/s_{R_1}, \kappa_{on_i} = \frac{k_{on_i} s_{R_1}}{{k^0_{R_1}}^2}, \kappa_{off_i} = \frac{k_{off_i}}{k^0_{R_1}}, \alpha_i = k^1_{R_i}/k^0_{R_1}, \beta_i = k_{ri}^1/k_r^0, \gamma = k_r^0/k^0_{R_1}, \delta = \frac{s_{R_2}}{s_{R_1}}, \epsilon = \frac{k^0_{R_2}}{k^0_{R_1}}$$

with $i = 1, 2$.

Here, $\sigma$ represents the synthesis rate constant of miRNA relative to that of the first RNA target. $\kappa_{on_i}$ and $\kappa_{off_i}$ represent miRNA binding and unbinding rates of the $i$-th RNA species. $\alpha_i$ describes the degradation rate of the $i$-th RNA in complex relative to that of its unbound form. $\beta_i$ is the degradation rate of miRNA in the $i$-th complex relative to its unbound form. $\gamma$ is the degradation rate of unbound miRNA relative to that of the first RNA species. $\delta$ and $\epsilon$ describe respectively the synthesis and the degradation rate of the second target with respect to the first.

#### 4. Estimation of relative target parameters in model M2

For parameters $\sigma$, $\kappa_{on_i}$, $\kappa_{off_i}$, $\alpha_i$, $\beta_i$ and $\gamma$ we adopted ranges estimated for the M1 model. For the remaining parameters $\delta$ and $\epsilon$ we chose the range $[10^{-1}, 10^{1}]$, to explore scenarios where one of the two targets is characterized by slower or faster synthesis and/or degradation kinetics with respect to the other. Parameter ranges for the nondimensional M2 model are reported in the following table, where $i = 1, 2$:

| **Nondimensional parameter** | **Biological meaning** | **Range** |
|---|---|---|
| $\sigma$ | Scaled miRNA synthesis rate | $[10^{-1}, 10^{1}]$ |
| $\kappa_{on_i}$ | Scaled binding rate | $[6, 6 \times 10^{3}]$ |
| $\kappa_{off_i}$ | Scaled unbinding rate | $[7.7 \times 10^{-2}, 7.7 \times 10^{1}]$ |
| $\alpha_i$ | Bound relative to unbound RNA degradation rate | [1/8, 16] |
| $\beta_i$ | Bound relative to unbound miRNA degradation rate | [1/8, 16] |
| $\gamma$ | Scaled miRNA degradation rate | $[10^{-1}/4, 10/4]$ |
| $\delta$ | Relative target synthesis rate | $[10^{-1}, 10^{1}]$ |
| $\epsilon$ | Relative target degradation rate | $[10^{-1}, 10^{1}]$ |

Table 6: Scaled M2 model's parameter ranges.

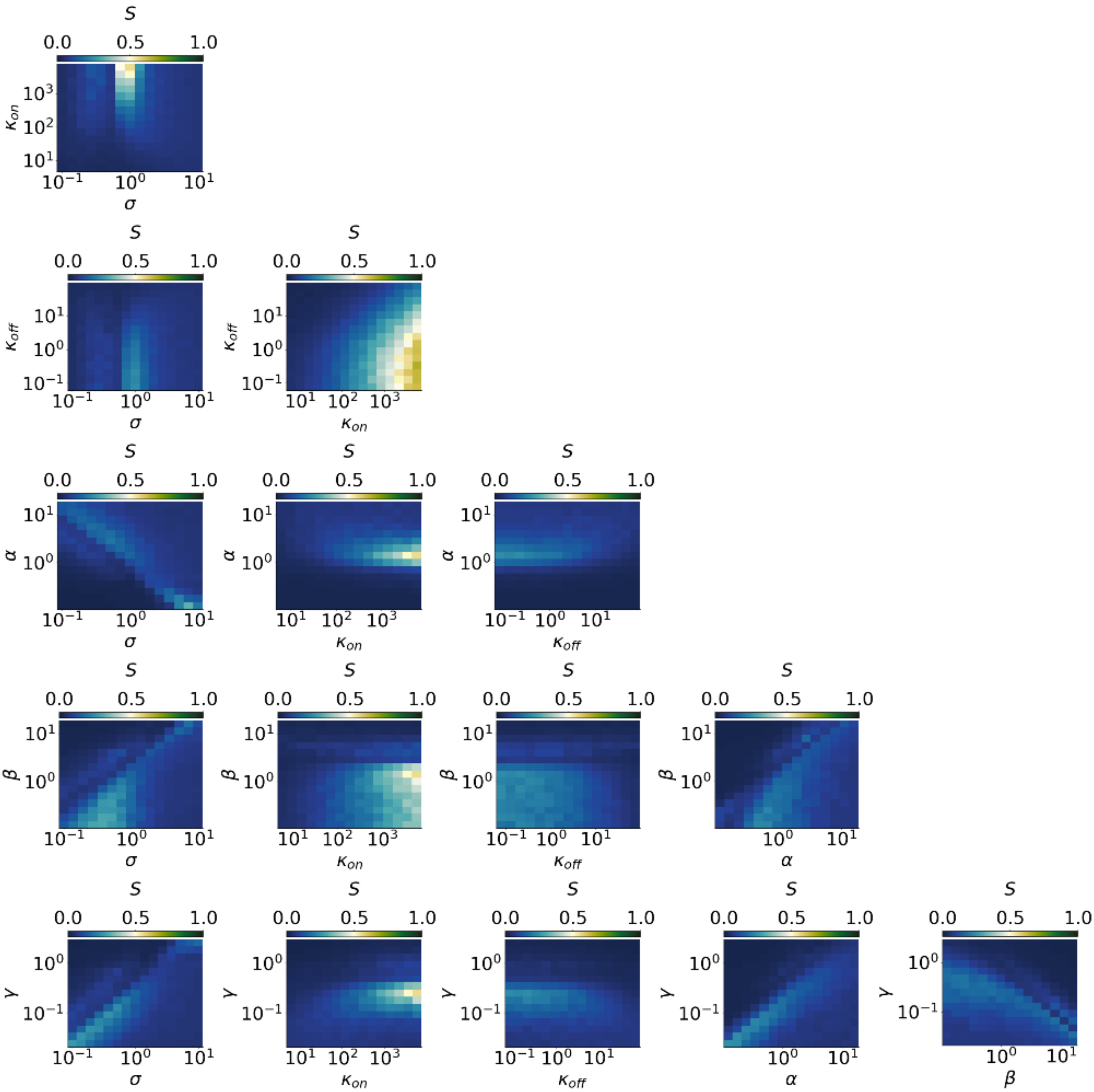




Figure 6: Selectivity computed as a function of parameter pairs of model M1. Each plot represents $S$ as a function of parameter values for a parameter pair of model M1.

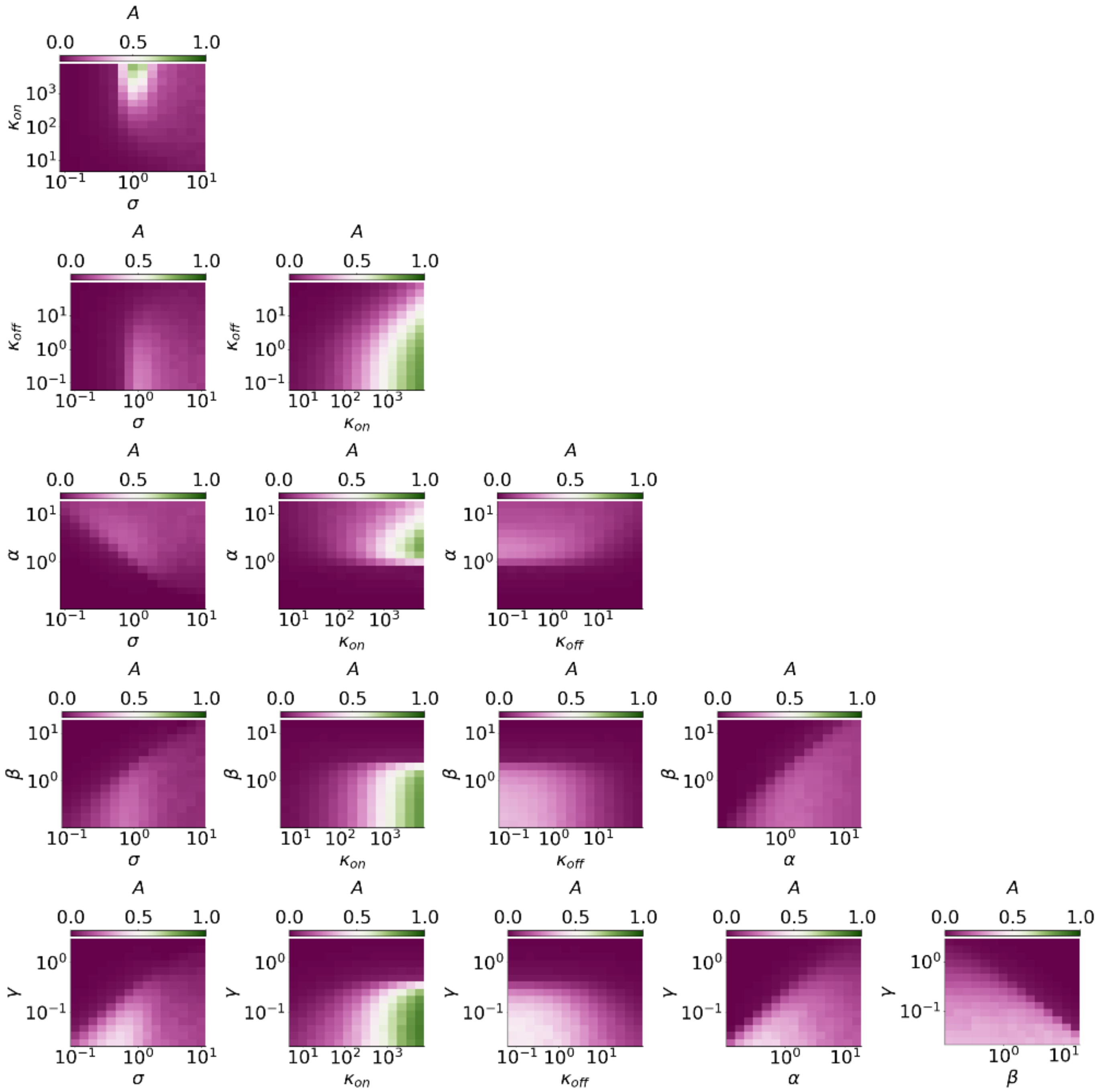




Figure 7: Advantage computed as a function of parameter pairs of model M1. Each plot represents $A$ as a function of parameter values for a parameter pair of model M1.

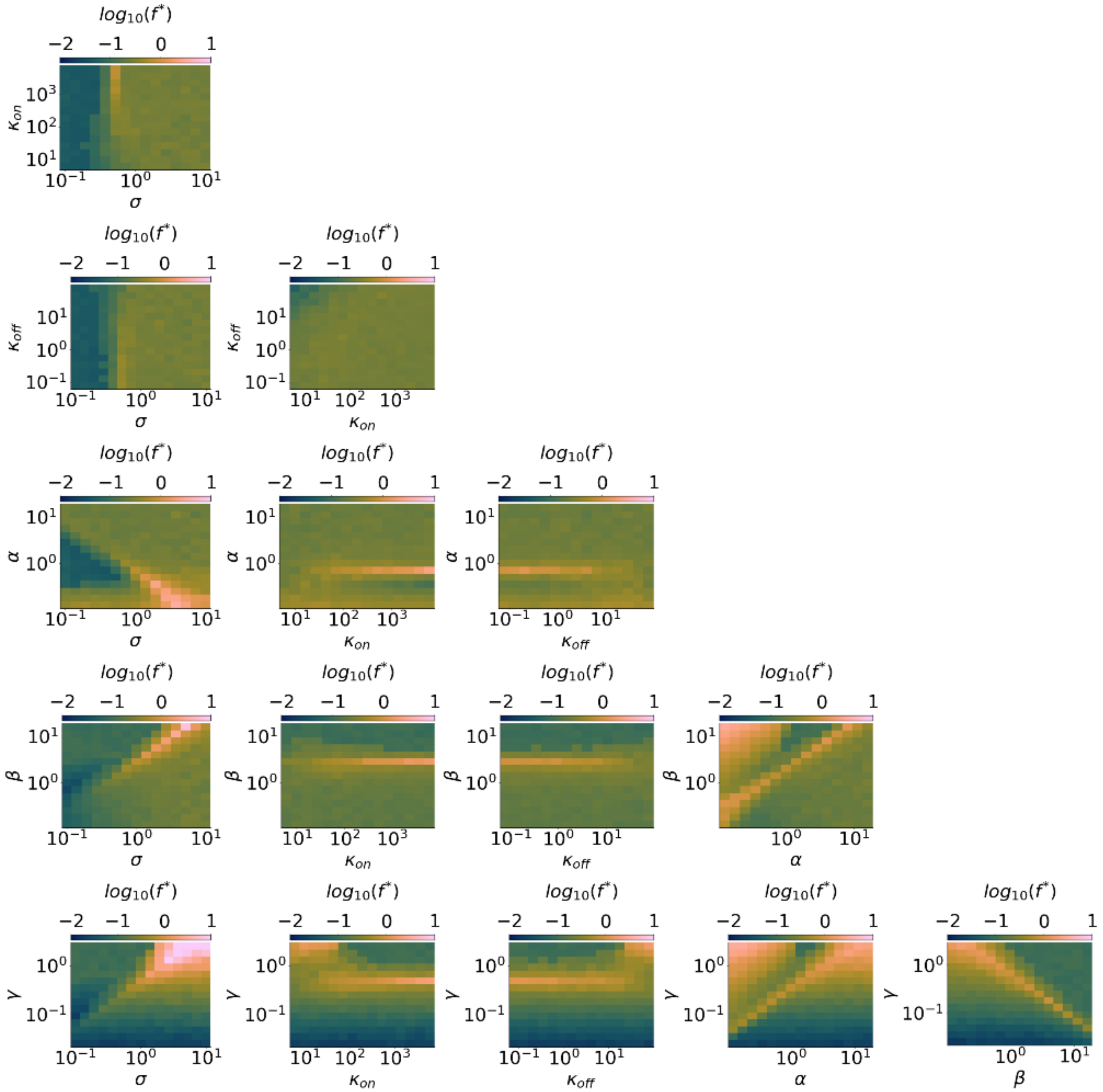




Figure 8: Preferred frequency computed as a function of parameter pairs of model M1. Each plot represents $f^*$ as a function of parameter values for a parameter pair of model M1.

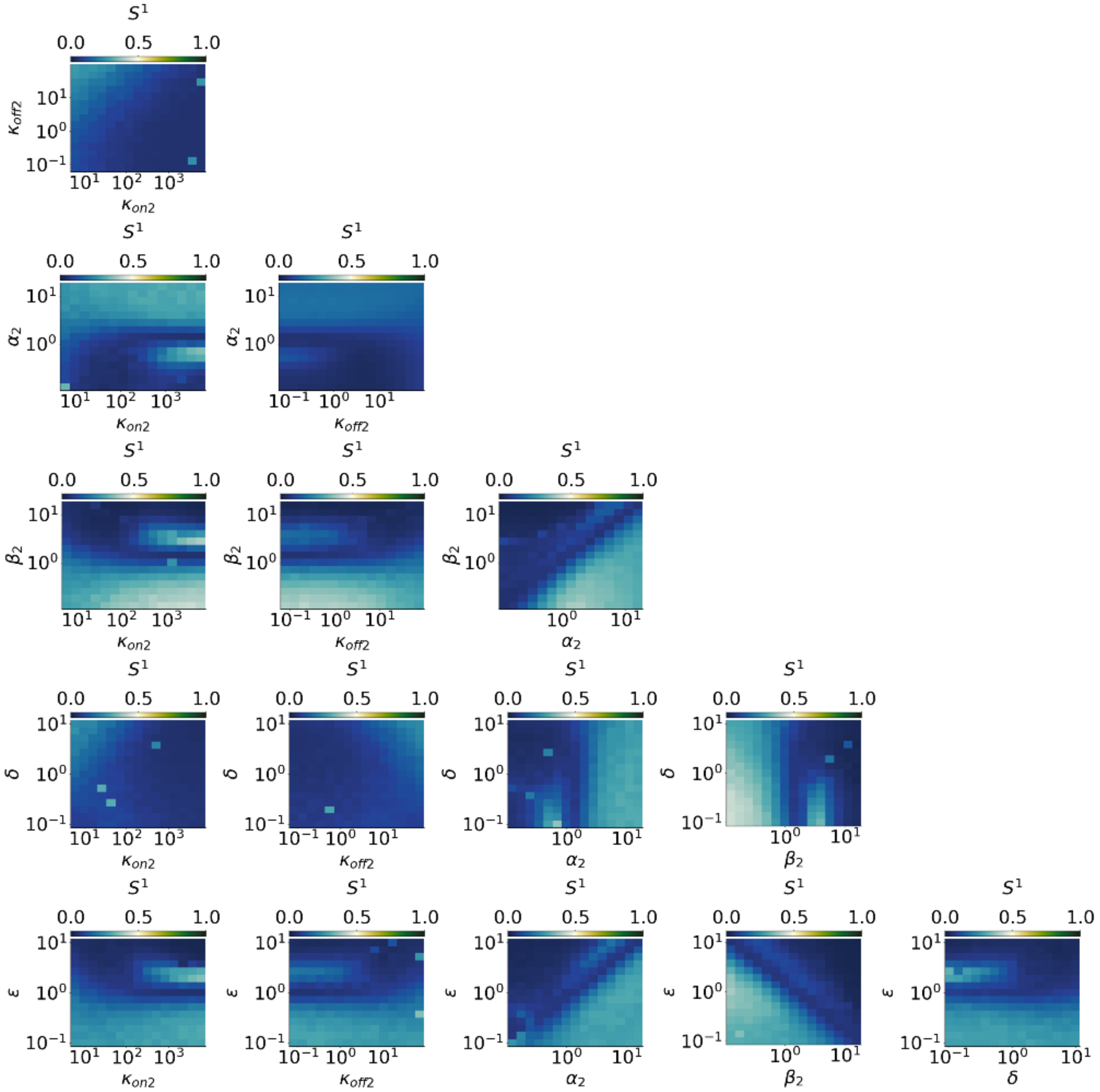




Figure 9: Selectivity of target $R^1$ computed as a function of parameter pairs relative to target $R^2$ in model M2. Each plot represents $S^1$ as a function of parameter values for a parameter pair of target $R^2$ in model M2.

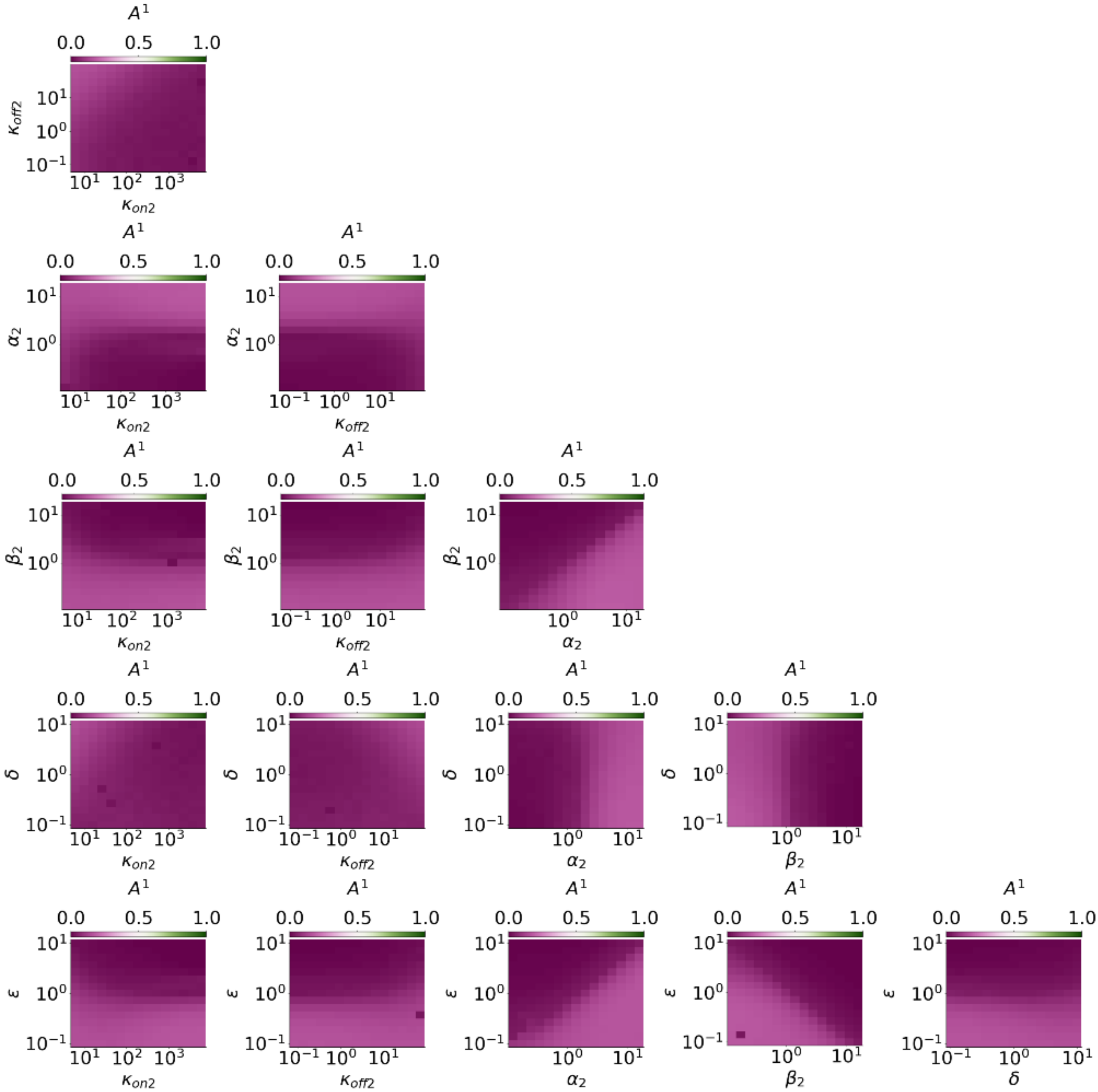




Figure 10: Advantage of target $R^1$ computed as a function of parameter pairs relative to target $R^2$ in model M2. Each plot represents $A^1$ as a function of parameter values for a parameter pair of target $R^2$ in model M2.

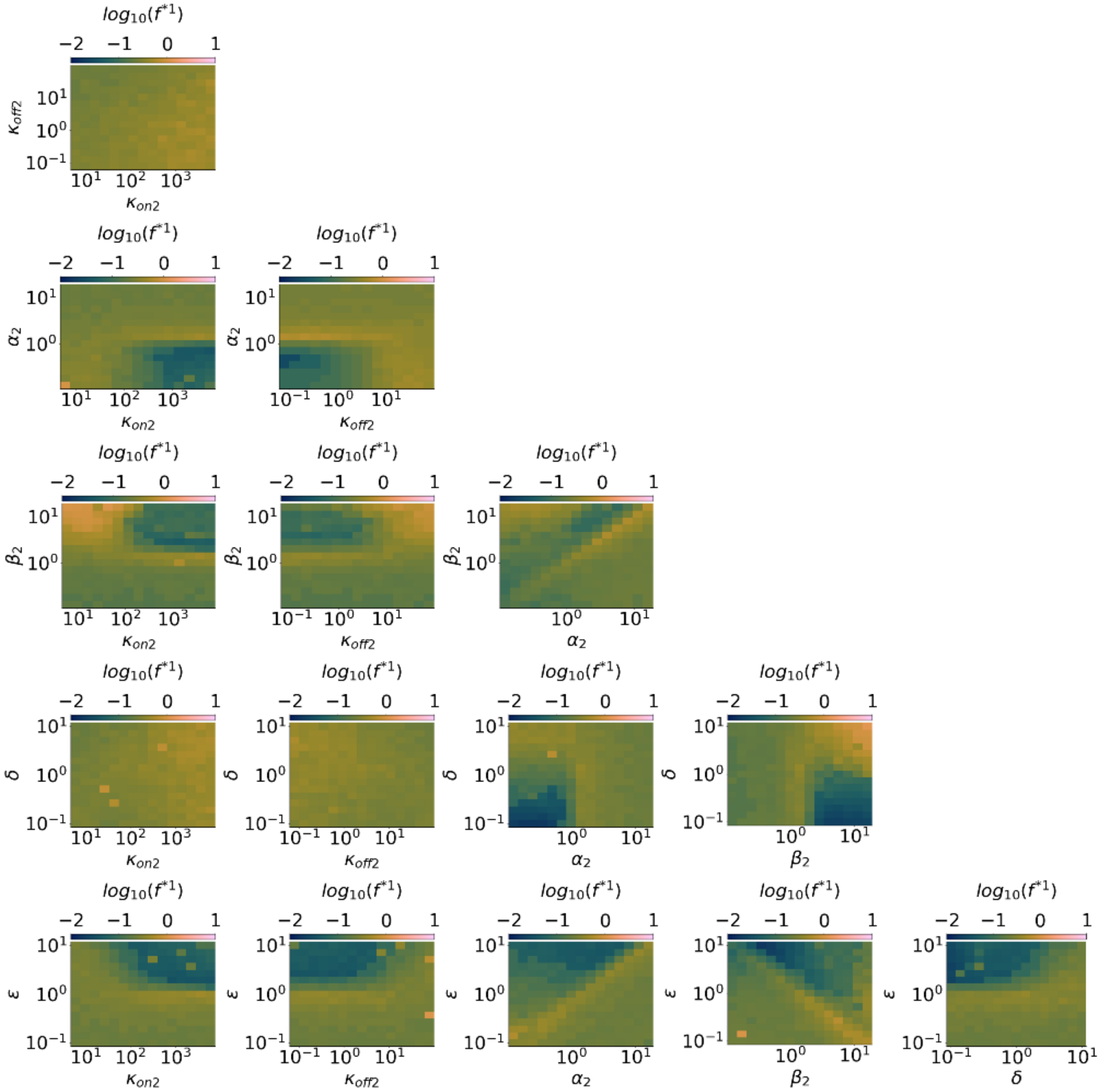


Figure 11: Preferred frequency of target $R^1$ computed as a function of parameter pairs relative to target $R^2$ in model M2. Each plot represents $f^{*1}$ as a function of parameter values for a parameter pair of target $R^2$ in model M2.

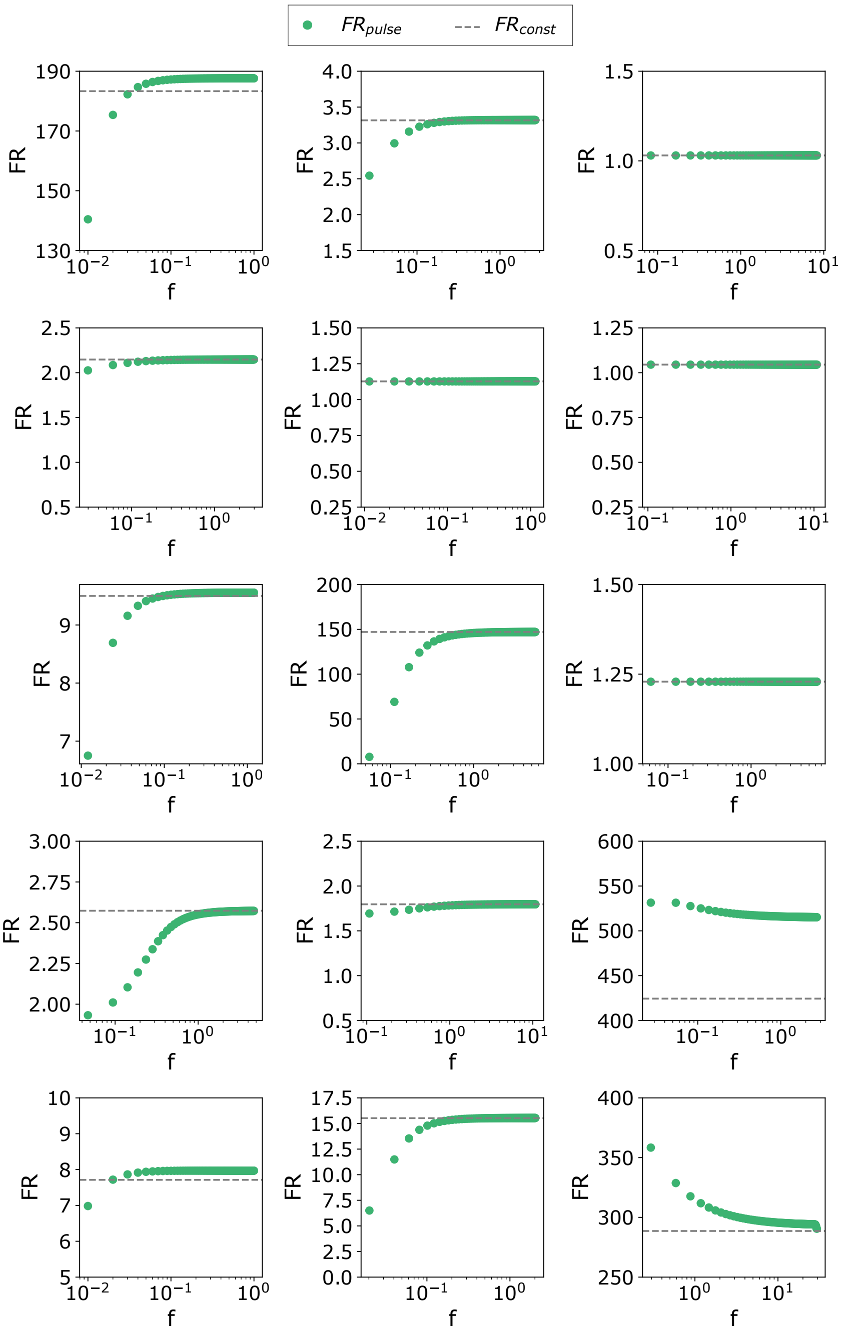


Figure 12: Fold repression given by constant input miRNA synthesis at the steady state. Each plot represents $FR_{const}$ (gray dashed line) and $FR_{pulse}(f)$ (green dots) computed in the timespan $[\tau, 2\tau]$ for a randomly sampled parameter set of model M1.
